# Synthetic Mimetics of Exosomal Lipid Profile Enhance Gemcitabine Delivery in Pancreatic Cancer

**DOI:** 10.1101/2025.10.24.684306

**Authors:** Tina Napso Shogan, Avi Schroeder, Lana Ginini, Malak Amer, Neta Regev-Rudzki, Ori Avinoam, Haguy Wolfenson, Ziv Gil

**Affiliations:** The Rappaport Faculty of Medicine and Research Institute, Technion - Israel Institute of Technology, Haifa 31096, Israel; The Luis Family Laboratory for Targeted Drug Delivery and Personalized Medicine Technologies, Department of Chemical Engineering, Technion, Haifa 32000, Israel; Department of Orthodontic and Craniofacial Anomalies, Rambam Health Care Campus, Haifa 3109601, Israel; Department of Biomolecular Sciences, Weizmann Institute of Science, Rehovot 7610001, Israel; Head and Neck Institute, The Holy Family Hospital, Nazareth 1641100, Israel

**Keywords:** Pancreatic cancer, Exosomes, Drug delivery, Lipid composition, Synthetic exosome mimetics, Phosphatidylserine

## Abstract

Pancreatic cancer ranks fourth among cancer-related deaths. Despite decades of research, the cure rate of disease remains disappointingly low. This dismal prognosis is due to late detection and to resistance of tumors to all known systemic therapies. Here, we build upon our previous findings that highlighted the selective uptake of tumor-associated macrophage-derived exosomes by cancer cells. We hypothesize that the prominent lipid contents on the exosome surface play a crucial role in facilitating selective cellular interactions, offering a novel avenue for developing efficient drug delivery platforms for pancreatic cancer. Lipids’ affinity toward cancer cells was measured, following mass spectral lipidomic analysis for screening the lipidic composition of macrophages and their derived exosomes. Subsequently, optimized and characterized synthetic exosomes underwent detailed intracellular trafficking studies in human pancreatic cancer cells, demonstrating enhanced cellular uptake kinetics. Notably, gemcitabine-loaded synthetic exosomes exhibited superior efficacy in inducing programmed cell death compared to both the free drug and conventional liposome formulations. The biodistribution examination of these synthetic exosomes underscored their potential for tumor specificity. *In vivo* experiments further demonstrated that the treatment with gemcitabine-incorporated synthetic exosomes inhibited tumor growth by nearly 50% compared to the administration of the free drug, indicating a significantly enhanced treatment efficacy. Our findings indicate that leveraging the lipid membrane composition of exosomes could lead to breakthroughs in drug delivery efficiency. This innovative strategy offers a promising direction in the ongoing battle against pancreatic cancer, highlighting the potential of exosomal lipid profiles for cancer therapy.

## Introduction

Pancreatic cancer stands as the third leading cause of cancer-related mortality in the United States [1], with projections indicating that it may ascend to become the second most significant cause of cancer-related death by 2030, following only lung cancer [2]. The challenging landscape of pancreatic cancer is marked by elusive early detection, rapid progression, high recurrence, emergence of drug resistance, and a deficiency in effective therapeutic approaches [3]. These factors collectively underlie the critical urgency to devise novel therapeutic and diagnostic strategies capable of alleviating the dismal prognosis of pancreatic cancer patients.

The complexity of the tumor microenvironment includes local hypoxia, chemokines, cytokines, and other secreted factors, resulting in the recruitment and differentiation of monocytic cells into tumor-associated macrophages (TAMs) [4,5]. During the advanced stages of pancreatic cancer, TAMs undergo a shift towards the M2 immunosuppressive phenotype, promoting processes such as tumorigenesis, lymphatic metastasis, and chemotherapy resistance, ultimately leading to bleak prognostic outcomes [6–8]. Significantly, the exosomes derived from M2 polarized-TAMs have a pivotal role in tumor progression, invasion, metastasis, and even chemotherapy resistance [9–11].

Within the context of cancer, exosomes derived from diverse cells in the tumor microenvironment influence cancer-associated processes such as cellular proliferation, invasion, and metastasis [12–14]. Exhibiting diameters ranging from 50 to 200 nanometers, exosomes are extracellular vehicles (EVs) present in the extracellular space and various body fluids. Filled with nucleic acids, proteins, and lipids, exosomes exhibit dynamic compositions that vary based on their origin cell types and the prevailing physiological and pathological conditions [15]. Their capability for molecular exchange between cells positions them as key mediators of intercellular communication [16], a quality that has triggered considerable interest in their potential applications for diagnostics and therapeutics.

Due to the unique lipid arrangement of the exosome membrane, which differs from the composition of their parent cell’s plasma membrane [15], lipids are recognized as potential clinical biomarkers for various diseases, particularly cancer. Similar to their role in cells, lipids are the building blocks of exosome membranes, contributing to their architecture, stability, and governing surface curvature [17]. These qualities provide exosome membranes with the essential stability to safeguard cargos from the risk of degradation during their transport within extracellular environments and bodily fluids. However, lipids abounding in the membrane of exosomes also function as bioactive molecules and play pivotal roles not only in vesicle structuring and stabilization but also in exosome biogenesis, secretion, internalization, and interactions with recipient cells [18].

In previous work, we unveiled the preferential internalization of exosomes derived from M2 macrophages into cancer cells within the heterogeneous tumor microenvironment [10]. This discovery sparked our interest in the unique components of the exosome membrane. Specifically, in this paper, we are focusing on the lipid composition of exosomes, examining their prominent lipid contents, and exploring their potential roles in facilitating these selective cellular interactions. We hypothesize that harnessing this advantage could pave the way for the development of synthetic exosome mimetics—a potential tool for efficient drug delivery in the battle against pancreatic cancer.

## Materials and Methods

### Liposomes Preparation

For HSPC formulation, Liposomes were composed of 65% mol hydrogenated soybean phosphatidylcholine (HSPC; Lipoid, Ludwigshafen, Germany); 5% mol N-(carbonyl-methoxypolyethylene glycol-2000)-1,2-distearoyl-sn-glycero-3-phosphoethanolamine (DSPE PEG 2000; Lipoid, Ludwigshafen, Germany); and 30% mol cholesterol (Sigma-Aldrich, Rehovot, Israel). The lipids were dissolved in absolute ethanol and warmed to 70°C. The lipid mixture was injected to 70°C warned 250mM ammonium sulfate. Additionally, PS formulation was composed of 10% mol 1,2-dioleoyl-sn-glycero-3-phospho-L-serine (18:1 PS, Lipoid, Ludwigshafen, Germany); 55% mol hydrogenated soybean phosphatidylcholine; 5% mol polyethylene glycol distearoyl-phosphoethanolamine and 30% mol cholesterol. The lipids were dissolved in chloroform and a thin lipid film was obtained by removing the solvent with rotary evaporator (BUCHI Labortechnik AG, Postfach, Switzerland) at 55°C. The film was hydrated with 250 mM ammonium sulfate. Both formulations were downsized by stepwise extrusion, using 400, 200, 100 and 80nm pore-size polycarbonate membranes (GE Healthcare Life Science, Whatman, Newton, Massachusetts, USA) in a LIPEX Extruder (Northern Lipids, Vancouver, Canada), for gaining homogeneous 100nm liposomes. Both liposomes’ solutions were dialyzed against 5% D(+)-glucose monohydrate solution (>1:250 volume ratio) using a 12-14 kDa dialysis membrane (Spectrum Laboratories Inc., USA) at 4°C and exchanged three times (after 1 hour, 4 hours, and overnight). Gemcitabine-encapsulated liposomes preparation was made using active loading method [19]. Gemcitabine hydrochloride (Sigma-Aldrich, Rehovot, Israel) was dissolved in 10% sucrose and was added to the ammonium sulfate liposomes at a final concentration of 1 mg/ml. the liposomes-gemcitabine mixture was agitated at 750 rpm for 1 hr at 70°C. The non-encapsulated drug was removed by dialysis, as described. Liposomes size (nm) and size distribution dispersity index (PDI) were measures by dynamic light scattering using Zetasizer Nano ZSP (Malvern Panalytical, Worcestershire, UK). Zeta potential and particles concentration (particles/ml) measurements were preformed using a Zetasizer Ultra (Malvern Panalytical, Worcestershire, UK). The liposomes were storage in 4°C not more than five weeks.

### Rhodamine-labelled Liposomes Preparation

Fluorescent liposomes were prepared using 16:0 Liss Rhodamine PE (Avanti Polar Lipids, Alabaster, AL, USA). Each of the formulations (HSPC and PS) were prepared as described above, after addition of 16:0 Liss Rhod PE to the lipid mixtures in molar percentage of 0.06%. The HSPC-lipid mixture was injected into phosphate-buffered saline (DPBS; Sartorius, Biological Industries, Beit Haemek, Israel), while thin lipid film was made from PS-lipid mixture. Then, the lipid film was hydrated with PBS.

For liposomal uptake compression experiments, the same molar ratio of 16:0 Liss Rhod PE was used for all formulations, while DSPE PEG wasn’t included. In short, HSPC liposomes were prepared in molar ratio of 70:30 HSPC: cholesterol. As for the other formulation, each one of the different lipids [1,2-dioleoyl-sn-glycero-3-phospho-L-serine (PS), L-α-phosphatidylinositol (PI), L-α-lysophosphatidylcholine (LPC), sphingomyelin (SM), Ceramide (Cer), and phosphatidylethanolamine (PE); Avanti] was added to the basic HSPC: cholesterol lipid mixture in molar percentage of 5% (or 10% for PS) at an expense of HSPC. LPC liposomes were made by adding to the HSPC: cholesterol lipid mixture only 0.5% mol in order to get the same liposomes size. In addition, PI lipid mixture was made by ethanol/DMSO (1:1, v/v). All formulations were prepared using ethanol injection despite the PS formulation, which were dissolved by chloroform and thin lipid film was performed. Hydrations were made using PBS and all liposomes were extruded as described above.

### Evaluation of Encapsuled Gemcitabine Concentrations

High-performance liquid chromatography (HPLC) device (1260 infinity, Agilent Technologies, Santa Clara, California, USA) equipped with Luna C-18 column (150 × 4.6 mm; Phenomenex LTP, Aschaffenburg, Germany) and ultraviolet detectors was used to quantify the encapsulated gemcitabine concentration. For preparing the samples for HPLC measurement, the gemcitabine encapsulated within the liposomes was extracted by mixing the liposomes with the same volume of the chloroform: methanol mixture 90:10 v/v. Then, the mixture was centrifuged at 6000g for 5 min at 4°C. The upper phase was collected into HPLC vials and were diluted 1:25 with potassium phosphate buffer (pH 6.0). Absorption was measured at a wavelength of 267nm. The mobile phase consisted of potassium phosphate buffer and methanol in ratio of 90:10 respectively, with flow rate of 1 ml/min. Sample injection volume was 20μl and the run time was 6 min. Linear calibration curve was generated on the peak areas in the chromatograms and utilized to quantify the concentration of the analyzed samples. Loading efficiency was determined as the percentage of the detected loaded drug concentration from the total drug concentration which was used for Gem-liposome preparation.

### Cryogenic Transmission Electron Microscopy (Cryo-TEM)

A drop of HSPC or PS liposomes sample was placed on a carbon-coated perforated polymer film, supported by a 200 mesh TEM grid, held by tweezers. The drop was quickly plunged into liquid ethane at its freezing point (– 183°C), cooled by boiling liquid nitrogen. Cryo-specimens were examined in a T12 G2 cryo-dedicated TEM (Eindhoven, The Netherlands), operated at 120kV, using Gatan 626 (Gatan, Pleasanton, California, USA) cryo-holders, at Technion center for electron microscopy of Soft Matter, the Wolfson ddepartment of chemical engineering. Specimens were examined in the low-dose imaging mode (no more than 20 electrons per Å^2^). Images were acquired digitally by a US1000 cooled charge-coupled-device camera (Gatan, Pleasanton, California, USA).

### M2-Macrophages Production

Peritoneal macrophages were generated following previously established procedures [20]. In short, specific-pathogen-free C57BL/6 mice were intraperitoneally injected with Brewer thioglycolate broth. After five days, the mice were euthanized, and their peritoneal cavity was rinsed twice with cold PBS to harvest the peritoneal exudate cells. Subsequently, the cells were centrifuged in a refrigerated centrifuge, counted, and seeded in 100mm petri dishes for 2 hours to allow macrophages to adhere to the plastic surface. Nonadherent lymphocytes were gently washed away. To promote alternative M2 activation, 20 ng/ml murine IL-4 (Sigma-Aldrich, Rehovot, Israel) was added to the culture media for 48-hour stimulation. The M2 polarization was confirmed by CD206 and Arginase-1 markers using flow cytometry and RT-PCR analysis.

### Macrophages-derived exosomes Purification

Macrophage-derived-exosomes were isolated as previously described [21]. Briefly, macrophages were cultured in serum-depleted medium containing IL-4 for 48-72 hours. The exclusion of serum from the media was done to prevent contamination with serum-derived exosomes. The condition media was first centrifuged at 300 ×g followed by centrifugation at 10,000 ×g, for eliminating dead cells and debris. Finally, the resulting supernatant was subjected to ultracentrifugation at 100,000 ×g to pellet the exosomes. The exosome pellet was resuspended in PBS and stored at -80°C until further analysis.

### Lipidomic Analysis

Ultra performance liquid chromatography (UPLC) system (ACQUITY, Waters Corporation, Milford, Massachusetts, USA) combined to mass spectrometer (MS; Vion IMS QTof, Waters Corporation, Milford, Massachusetts, USA) were used for lipidomic analysis of M2 macrophages and their -derived exosomes. Chromatographic conditions were produced as described before with some adjustments [22]. Shortly, chromatographic separation was performed using ACQUITY UPLC BEH C8 column (1.7 μm, 2.1×100 mm; Waters Corporation, Milford, Massachusetts, USA), which was maintained at 40°C. the separation was achieved using two mobile phases. Mobile phase A consisted of a mixture of DDW: Acetonitrile: Isopropanol in a ratio of 46:38:16 (v/v/v) with 1% 1 M NH4Ac and 0.1% glacial acetic acid. On the other hand, mobile phase B was prepared with DDW: Acetonitrile: Isopropanol in a ratio of 1:69:30 (v/v/v) with 1% 1 M NH4Ac and 0.1% glacial acetic acid. The chromatographic system operated with a flow rate of 0.4 mL/min, and total run time was set to 25 minutes. The mobile phases produced linear gradient. Initially, mobile phase A was run at 100% for 1 minute, after which it was gradually reduced to 25% over the course of 11 minutes. This was followed by a further decrease to 0% for 4 minutes. Subsequently, mobile phase B was introduced and maintained at 100% for 5.5 minutes. Then, mobile phase A was briefly restored to 100% for 0.5 minutes. Finally, the column was equilibrated at 100% mobile phase A for 3 minutes. The MS parameters were configured as follows: the source and de-solvation temperatures were maintained at 120 °C and 450 °C, correspondingly. For positive ionization mode, the capillary voltage was set to 3.0 kV, while for negative ionization mode, it was set to 2 kV, with a cone voltage of 40 V. De-solvation gas (nitrogen) was used at a flow rate of 800 L/h, and the cone gas flow rate was set at 30 L/h. The MS operated in full scan high-definition resolution mode, covering a mass range of 50-2000 Da. For the high-energy scan function, a collision energy ramp of 20-80 eV was applied, while for the low-energy scan function, -4 eV was applied.

The LC-MS data was analyzed and processed using UNIFI software (Version 1.9.3, Waters Corporation, Massachusetts, USA). Lipid species identification was achieved by comparing accurate mass (<3 ppm), fragmentation pattern, retention time, and ion mobility values with an in-house lipid database. Subsequently, the peak intensities of the identified lipids were normalized to both the internal standards and the amount of tissue utilized for the analysis.

### Cell Cultures

BxPC-3 and MIA PaCa-2 were obtained from American Type Culture Collection (ATCC). KPC cells were generated in the lab from KPC mouse pancreatic tumor with KRAS G12D and TP53 R172H mutations, and pdx-1 CRE insertion [10,23–25]. KPC and BxPC-3 were cultures in RPMI-1640 medium (ATCC) and MIA PaCa-2 were cultured in Dulbecco’s modified Eagle medium (Sartorius, Biological Industries, Beit HaEmek, Israel). The mediums were contained 10% (v/v) fetal bovine serum (Sartorius, Biological Industries, Beit HaEmek, Israel), 1% (v/v) penicillin-streptomycin solution (100 IU/ml Penicillin G Sodium Salt and 100 µg/ml streptomycin sulfate) and 4mM L-glutamine (Sartorius, Biological Industries, Beit HaEmek, Israel). All cells were incubated at 37°C in a humidified atmosphere containing 5% CO2. Each cell line was split regularly when attaining 70–80% confluence. All cells were tested routinely for mycoplasma.

### Cellular uptake of liposomes

KPC, MIA PaCa-2, or BxPC-3 cells were seeded in a 12-well plate at respective densities of 10^5^, 2×10^5^ and 3×10^5^ cells per well. Following an overnight incubation, the culture medium was replaced with a fresh medium containing rhodamine-labelled liposomes at a concentration of 10^6^ liposomes per single cell. After a two-hour incubation, the cells were washed and stained with fixable viability dye (efluor 450, eBioscience, Thermo Fisher Scientific, Waltham, Massachusetts, USA). Finally, the cells were analyzed using BD High Throughput LSR Fortessa II Cell analyzer (BD Biosciences, San Jose, California, USA), and the acquired data were analyzed using FlowJo software (FlowJo, LLC, Ashland, Oregon, USA). The median rhodamine intensity was measured in the live cell population of each sample. To assess the variations in cellular uptake among the different liposome formulations, the median values obtained for each liposome treatment were normalized to the corresponding liposome fluorescence levels, as measured using a microplate reader (Infinite 200 PRO, TECAN, Männedorf, Switzerland).

### Endocytosis Examination

MIA PaCa-3 and BxPC-3 cells were seeded in 35 mm glass bottom dishes at a density of 2 × 10^5^ and 4 × 10^5^, correspondingly. Following a 24-hour incubation to ensure cell attachment, 1 mM PS-liposomes labelled with rhodamine were added into the culture media. Subsequently, the dishes were divided into two groups: one group was returned to the incubator for 37°C incubation, while the other group was placed in a 4°C. After two hours of liposome treatment, the cells were washed and fixed by 4%PFA. The nuclei were stained (Hoechst 33342, Sigma-Aldrich, Rehovot, Israel) and the cells were imaged by confocal microscopy (LSM700, Zeiss, Germany). The analysis was performed using ImageJ software (NIH), which involved normalizing the measured rhodamine intensity to the number of cells in each frame.

### Intercellular Liposomal Tracking

For lysosome colocalization experiments, 2 × 10^4^ MIA PaCa-2 cells were seeded in 35 mm glass bottom dishes with culture medium containing CellLight™ Lysosomes-GFP, BacMam 2.0 (Invitrogen, Thermo Fisher Scientific, Waltham, Massachusetts, USA) for overnight incubation. On the following day, the cells were washed, and the culture medium was refreshed. Subsequently, 1 mM PS-liposomes labelled with rhodamine were added into the fresh medium and incubated for two hours. At each experimental time point, the medium was replaced with phenol-free medium, and Hoechst 33342 (NucBlue Reagent; Invitrogen, Thermo Fisher Scientific, Waltham, Massachusetts, USA) was added for staining the cell nuclei. The cells were imaged by confocal microscopy (LSM700, Zeiss, Germany) with humified chamber.

To perform nucleus colocalization analysis, cells were seeded at the specified density as mentioned above. Once cell attachment was ensured, they were exposed to 1 mM PS-liposomes labelled with rhodamine for the desired duration. At each experimental time point, the cells were washed, and the culture medium was changed to phenol-free medium. Hoechst 33342 (NucBlue Reagent) was added for nucleus staining, and cells were imaged using confocal microscopy (LSM700, Zeiss, Germany) with a humidified chamber.

The colocalization analysis was made using Imaris software (Bitplane, Oxford Instruments, Belfast, UK). Z-stack confocal images were utilized to create 3D model images. The lysosome, nucleus, and liposome structures were reconstructed, and liposomes that were located at a distance of ≥0 from the lysosome or the nucleus were quantified. Then, the number of liposomes in this population was normalized to the volume of the respective organelle structures.

### Programmed Cell Death

Cleaved PARP1 expression was assessed using Western Blot. In short, 5 × 10^5^ MIA PaCa-2 and BxPC-3 cells were seeded in a 6-wells plate. After attachment, cells were treated for 24- and 48-hours with 1 μM free gemcitabine and liposomes containing the same concentration of drug. Cell lysates were produced using M-PER Mammalian Protein Extraction Reagent (Thermo Fisher Scientific, Waltham, Massachusetts, USA), and the protein concentrations were determined using the Bradford assay. Equal protein amounts were loaded onto SDS-PAGE gels. After gel transfer to nitrocellulose membranes, the membranes were blocked with 5% BSA for 1 hour. The primary antibody (anti-cleaved PARP1 antibody, Abcam ab32064) was applied and incubated overnight at 4°C, followed by the addition of the secondary antibody for 1 hour at room temperature. To compare the relative changes in protein expression among the experimental groups, the ratio of the target protein intensity to the housekeeping gene (β actin) intensity was calculated.

Caspase 3/7 expression was examined using fluorescent microscopy. MIA PaCa-2 and BxPC-3 cells were seeded in a µ-slide 8 well glass bottom (ibidi, Gräfelfing, Germany) at a density of 2 × 10^4^ and 1.5 × 10^4^, respectively. Subsequentially, the cells were treated with free gemcitabine and gemcitabine-loaded liposomes as described above. After the desired treatment duration, cells were incubated with Caspase-3/7 detection reagent (CellEvent, Invitrogen, Thermo Fisher Scientific, Waltham, Massachusetts, USA) for 1 hour. Then, cells were washed and fixed and the nuclei were stained by Hoechst 33342. Cells were imaged using fluorescent microscopy (Zeiss Axio observer Zeiss, Germany) and the images were analyzed using ImageJ software. To assess the nuclear expression of caspase 3/7, the nuclei in each image were classified. Then, the intensity of caspase 3/7 within the selected nuclear area was measured.

### Dose Response

Cell viability was assessed in order to produce dose-response curves. Briefly, BxPC-3 and MIA PaCa-2 cells were seeded in a 96-well plate at densities of 1.2 × 10^5^ and 5 × 10^4^, correspondently. The following day, the culture medium was replaced with fresh medium containing either gemcitabine loaded liposomes or free gemcitabine at gradually increasing concentration. After 24 or 48 hours, XTT solution (Sartorius, Biological Industries, Beit HaEmek, Israel) was added to each well. Following 1-2 hours incubation, the absorbances were measured using plate reader at a wavelength of 450-500 nm with a reference absorbance at 630-690 nm. The viability was calculated by the percentage of each treatment absorbance from the untreated cells total absorbance.

### Therapeutic Efficacy

The animal studies were in accordance with a protocol approved by the Institutional Animal Care and Use Committee (IACUC) of Technion. The animals’ well-being was monitored routinely with guidance of qualified veterinarians and was documented in scoring tables.

Three million MIA PaCa-2 cells suspended in 100 µl PBS were injected subcutaneously to the right flank of 8-week-old athymic nude mice. Four weeks later, once the tumors average size reached to 100 mm^3^, mice were divided to four treatment groups: (1) ctrl-control group, treated with 5% D(+)-glucose solution (n=4), (2) Lipo-Gem group-10mg/kg gemcitabine encapsulated in PS liposomes (n=6) , (3) Gem 10-free gemcitabine in concentration of 10mg/kg (n=5) and (4) Gem 100-free gemcitabine in concentration of 100mg/kg (n=6). All the treatments were administered intravenously once a week for three weeks. Mice weight and the tumor size were measured twice a week. Tumor volume was measured using a calliper and calculated as (width× width× length)/2. For tumor size comparisons in the different treatment groups, each mouse’s tumor size was normalized to the measured size at the first day of the experiment (first day of injection, prior the injection). The same was applied to compare the body weight of the mice. After sacrifice, the tumors were harvested and were added to formalin solution. The fixed tumors embedded in paraffin, sectioned, and stained for hematoxylin and eosin (H&E) for histology analysis. To assess apoptosis within the tumors, Terminal deoxynucleotidyl transferase dUTP nick end labelling (TUNEL) staining (ab206386; Abcam, Cambridge, Massachusetts, USA) was conducted following the manufacturer’s protocol. For this analysis, three sections from 3 to 4 tumors per treatment group were stained. From each section, fifteen random fields were captured, and the areas containing 3,3’-Diaminobenzidine (DAB) were quantified as a percentage of the total apoptotic area.

The quantification of gemcitabine release from the liposomes was performed using HPLC. The batch of gemcitabine encapsulated-liposomes which prepared for this experiment, was tested to determine the amount of gemcitabine retained inside the liposomes until the last administration, ensuring the delivery of encapsulated gemcitabine. Additionally, the size and PDI of the liposomes were monitored throughout the study to confirm their stability.

### Biodistribution

MIA PaCa-2 cells were injected into athymic nude mice, and the mice were monitored throughout the experiment as described above. Six weeks later, once the tumors average size reached 400 mm^3^, mice were divided into two groups. One group were injected intravenously with rhodamine-labelled Gd-loaded PS liposomes, while the other group was injected with HSPC liposomes. Each group included n=6 mice. Four hours after treatment, all mice were imaged using preclinical *in vivo* imaging system IVIS SpectrumCT (PerkinElmer, Massachusetts, USA) an excitation of 570 nm and emission of 620 nm for detecting accumulation of rhodamine-labelled liposomes. After mice were euthanized, all collected tissues (tumor, liver, lungs, heart, kidneys, pancreas, and brain) were imaged by IVIS with the same settings as detailed.

### Liver Function and Toxicity

Immediately prior to sacrifice, a blood sample of 500 µl was collected from the facial vein of both the control and liposomes groups. Blood serum was produced from the samples by allowing the blood to clot undisturbed at room temperature for 1 hour, followed centrifugation at 1500xg 4°C for 15 min. The serum samples were analyzed for alanine aminotransferase (ALT) and aspartate transaminase (AST) using Catalyst One, IDEXX Laboratories, Inc., Westbrook, Maine, USA.

The liver, lungs, heart, pancreas, spleen, and kidneys were extracted and immediately added to formalin solution. The tissues were fixated at least 24 hours before embedding in paraffin and sectioning. The different sections were stained for H&E according to the standard protocol.

### Statistical analysis

The data underwent initial normality testing, followed by the appropriate parametric or non-parametric statistical tests for each dataset. For normally distributed data, unpaired Student’s t-test, one-way ANOVA or two-way analysis of variance ANOVA were conducted as statistical analyses. Non-parametric testing was performed using the Kruskal-Wallis test. All statistical analysis and graph plotting were performed using Prism GraphPad software (version 10.1.2). Statistical significance was determined at a threshold of P < 0.05, with significance levels represented as ***p < 0.0001, ***p < 0.001, **p < 0.01, *p < 0.05, and ns indicating non-significance.

## Results and Discussion

### Identification of Exosomes’ Lipidic Surface Profile and Its Contribution to Cellular Uptake

Like the cell plasma membrane, exosome membranes are composed of cholesterol, sphingolipids, and different glycerophospholipids. However, the lipid composition of exosome membranes differs from that of the membranes of their parent cells. Multiple studies have characterized the lipid composition of exosomes originating from various cells (extensively reviewed by Skotland *et al*. [26,27]). In general, exosomes display higher levels of cholesterol, sphingomyelin (SM), phosphatidylserine (PS), and glycosphingolipids compared to their parent cells. Conversely, the abundance of phosphatidylcholine (PC) and phosphatidylinositol (PI) is higher in the cells themselves than in their derived exosomes. Notably, the content of phosphatidylethanolamine (PE) remains consistent in both the parent cells and exosomes.

As mentioned, we previously showed that macrophages derived exosomes (MDEs) are selectively internalized by pancreatic cancer cells [10]. To study the contribution of exosomes’ lipid composition in their communication with cancer cells, we focused on the lipid profile of exosomes derived from M2-polarized murine peritoneal macrophages. In pancreatic cancer, there is a predominant skewing of macrophages in the tumor microenvironment towards M2 polarization state. This polarization pattern promotes an anti-inflammatory phenotype, contributing to the development of immunosuppressive and pro-tumorigenic characteristics [6]. We performed lipidomic analysis on MDEs and their parent cells-M2 TAMs (Figure S1). Approximately 210 lipid species across various families were identified using liquid chromatography-mass spectrometry (LC-MS) analysis, encompassing phospholipids, glycolipids, glycerolipids, and free fatty acids. Notably, cholesterol was excluded from this analysis. For comparative purposes, lipid intensities identified in both cells and derived exosomes were normalized against internal standards and protein concentrations. Subsequently, the relative abundance of each lipid species was calculated as a percentage of the total lipidome detected.

Next, in this study we predominantly focused on phospholipids, including sphingolipids and glycerophospholipids, due to their dominance in the lipidic composition—constituting 80% and 90% of the total lipidome in exosomes and cells, respectively (Figure S1A). In particular, our analysis concentrated on various phospholipid classes present in both exosomes and macrophages, namely SM, PC, PE, ceramide (Cer), PS, and PI (Figure S1B).

We observed that numerous saturated phospholipid species, prevalent in cells, were completely absent in exosomes, aligning with previous findings [27]. Notably, two phosphatidylglycerol (PG) species detected in cells were entirely undetectable in exosomes. Moreover, lysophosphatidylcholine (LPC), a monoradyl derivative of PC, was found to be significantly more abundant in exosomes compared to cells (Figure S1C).

It is crucial to recognize that the identification of lipid species in this study was constrained by the scope of the lipid library used during analysis. As a result, there is a possibility that certain species not represented in the library were not detected. This limitation may have implications for the comprehensiveness of our findings and the subsequent interpretations derived from it.

Considering both the existing knowledge about exosomes lipid composition [26] and our lipidomic analysis, we highlighted six lipids for investigating the role of the different lipids in cellular internalization of exosomes. PE, PI, PS, SM, LPC and Cer were recruited to evaluate the contribution of each one of them to cellular uptake, in the mission to design an effective drug delivery platform, based on exosome lipid composition. To assess the input of each lipid class on cellular uptake, we fabricated liposomes with each one of the listed lipid classes. The reference liposomes, which we called “basic” or simply HSPC, were composed of HSPC and cholesterol (70:30 %mole). Generally, PC and cholesterol both stand out as predominant lipids in membranes [28]. The candidate lipids were added into the basic formulation in concentration of 5% or 10% at the expense of HSPC, except LPC which was 0.5%. The liposomal tool allowed to preserve the physical properties of exosomes, as they also have lipid bilayer membrane, providing a possibility for encapsulating cargos and to conserve the nanometric size (Table S1).

To quantify the cellular uptake, we incubated KPC and MIA PaCa-2 cells with rhodamine-labelled liposomes for two hours and analysed the internalization by flow cytometry (Figure 1). Recruitment of PS into the basic HSPC liposomes extremely elevated cellular uptake of the liposomes in both KPC and MIA PaCa-2 cells, compared to the unmodified HSPC liposomes. In contrast, the addition of all other lipids had either negligible or minimal contribution to the uptake efficacy to that of HSPC-liposomes. The powerful effect of PS liposomes raised the question whether increasing the percentage of PS would improve the uptake. Indeed, 10% PS liposomes displayed significantly higher uptake by both examined cell lines, compared to the lower percentage PS formulation.

**Figure 1:**
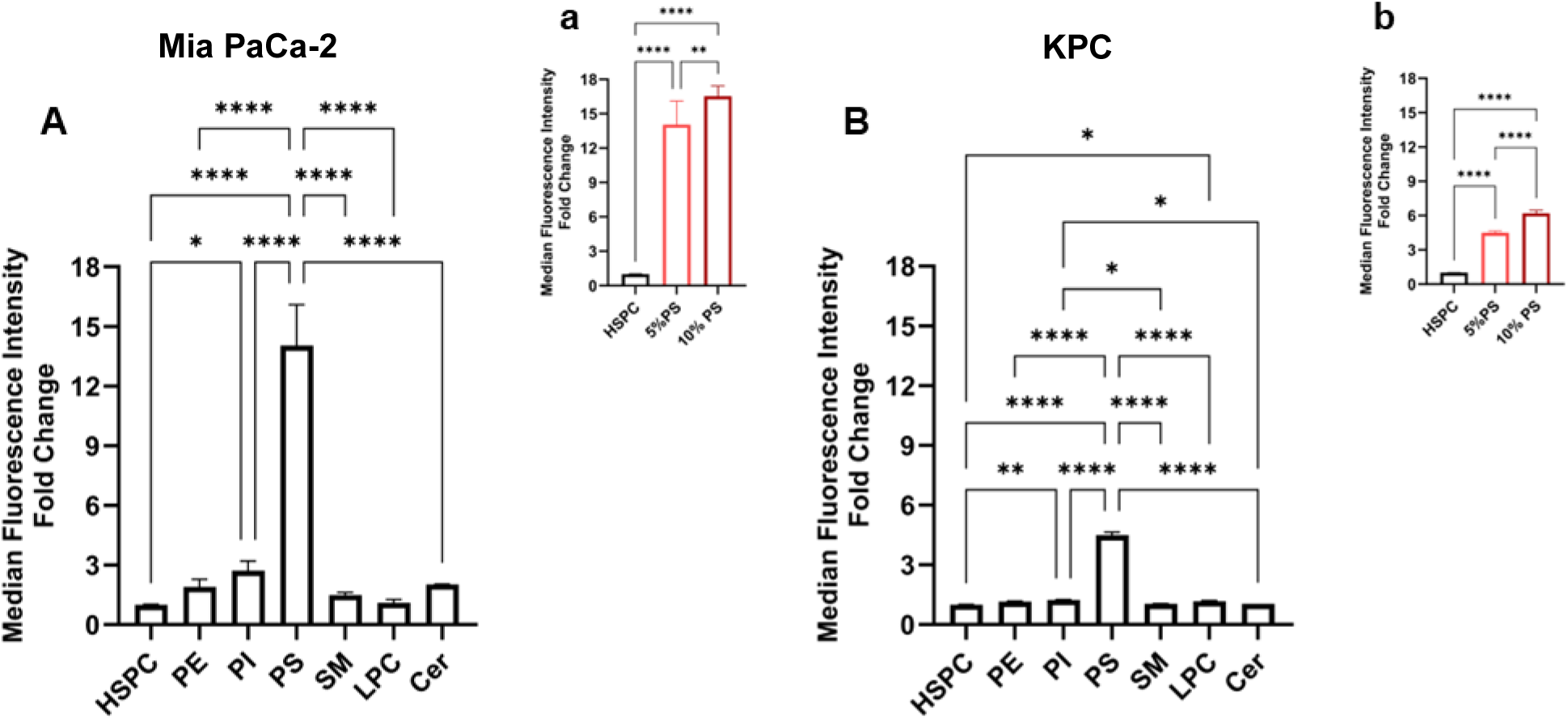
The impact of incorporating various exosomal lipids into liposomes on uptake by pancreatic cancer cells. Mia PaCa-2 **(A,a)** and KPC **(B,b)** cells were treated with rhodamine-labelled liposomes for two hours at identical particle concentration. The contribution of each exosomal lipid to cellular uptake was assessed using flow cytometry. The median intensity values were normalized to those of HSPC. The data is presented as mean of ± SD (n=3), statistical analysis was perfumed using through one-way ANOVA followed by subsequent multiple comparisons test. Significance levels: *\*p* < 0.05, *\*\*p* < 0.01, *\*\*\*\*p* < 0.0001, any other unpresented comparison was not significant with *p* > 0.05.

These findings align those of Lu *et al*., who developed exosome-mimicking liposomes (formulated with DOPC/SM/Chol/DOPS/DOPE 21/17.5/30/14/17.5, mol/mol) for siRNA delivery, achieving significant cellular uptake and silencing efficacy [29]. Similarly, the efforts of Sakai-Kato *et al*. to design liposomes that mimic the lipid composition of HepG2-derived exosomes further validate the critical role of lipid composition, particularly the impact of DOPS, in enhancing cellular internalization [30]. Together, these studies support our findings and underscore the significance of the incorporation of PS in cellular uptake.

### Optimization of The Liposomes

To adapt the 10% PS liposomes for potential drug delivery in pancreatic cancer treatment, we incorporated polyethylene glycol (PEG) into the formulation. Given that PEG is a crucial component that enhances stability and bioavailability of the liposomes, its addition was necessary to optimize the liposome formulations for this application [31]. The PEGylated HSPC and PS liposomes were characterized by their size, polydispersity index (PDI), particle concentration and zeta potential (Figure 2A-D, Table S2, Figure S2). Moreover, we examined the particles morphology using cryo-TEM, in which the spherical shape of the particles was noticeable, and the uniform dispersity of particle population was verified additional to the measured PDI (Figure 2G). We also validated the DLS measured size by measuring the particles diameter in the cryo-TEM images (Table S2). Storage stability was examined by measuring the size and PDI of the particles once a week. Throughout the examined period, both HSPC and PS formulations remained remarkably stable and homogenous (Figure 2H-I).

**Figure 2:**
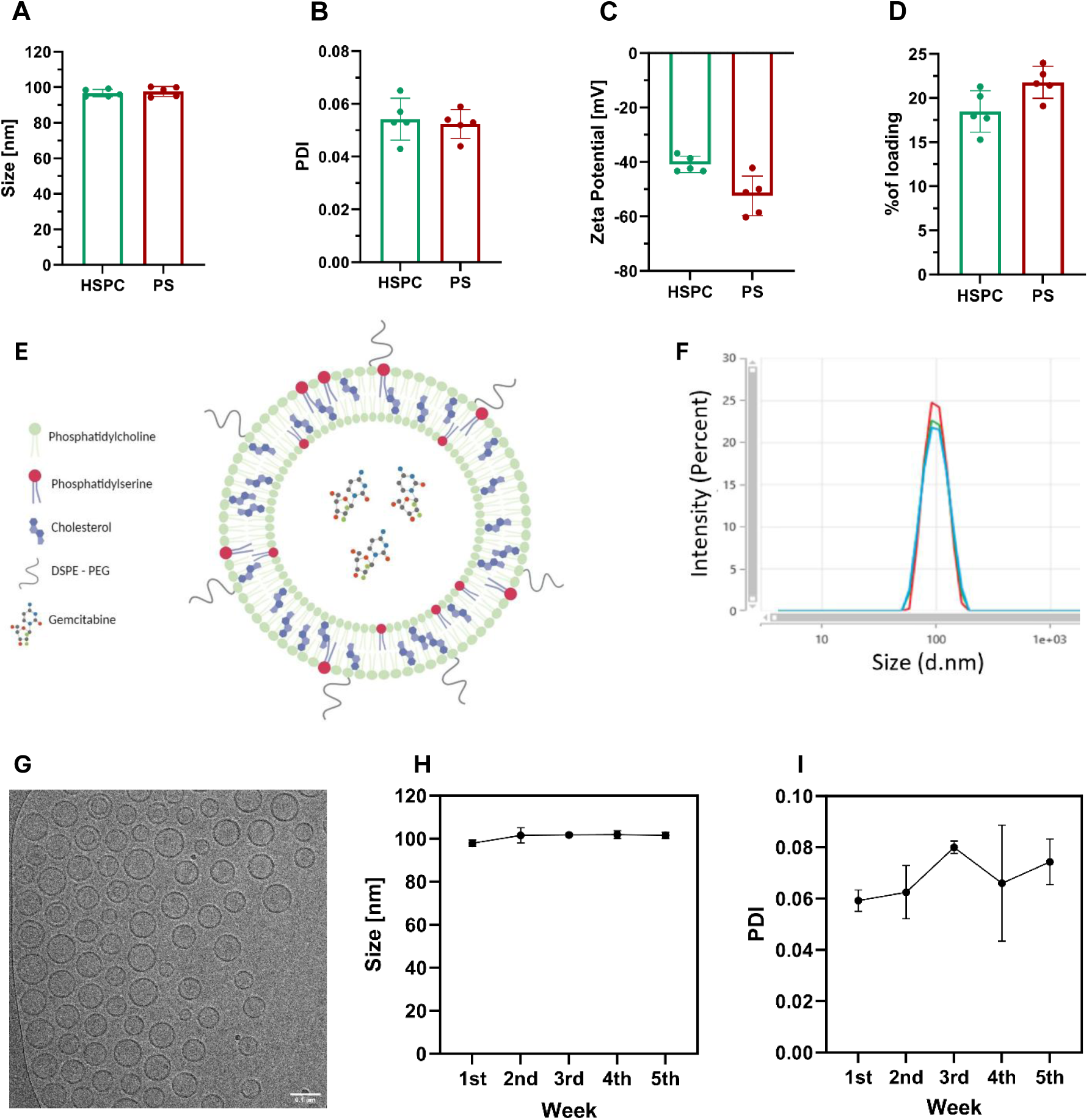
Characterization of PEGylated PS liposome. **(A)** The size, **(B)** polydispersity index (PDI), **(C)** particle concentration and **(D)** zeta potential of PEGylated HSPC and PS liposomes were evaluated using dynamic light scattering. The data is presented as mean ± SD (n=5). **(E)** Schematic illustration of 100 nm liposomes composed of HSPC (hydrogenated soybean phosphatidylcholine), cholesterol, PS (,2-dioleoyl-sn-glycero-3-phospho-L-serine ) , and DSPE– PEG-2000 (polyethylene glycol distearoyl-phosphoethanolamine), 55:30:10:5 molar ratio, loaded with gemcitabine (created by BioReander.com); **(F)** PS liposomes’ size was measured by dynamic light scattering; **(G)** Liposomes’ spherical shape and their uniform particle distribution were observed by cryo-TEM imaging, scale bar, 0.1 μm; The stability of PS liposomes under storage conditions (4°C) was analyzed by size **(H)** and PDI **(I)**. The data is presented as the mean ± SD (n=3).

Next, Gemcitabine, a long-standing drug for treating pancreatic cancer, was loaded into the liposomes via the active loading method (Figure S3). Typically, weak bases with a pKa of ≤ 11 and a log P value between −2.5 and 2.0 are considered suitable candidates for active loading [32]. Despite gemcitabine fulfilling these criteria, with pKa = 3.6 and logP = -1.5, achieving efficient encapsulation into liposomes via active loading method remains limited [33–35]. Efforts to enhance drug encapsulation efficiency have been documented [34,35]; some of these methodologies were tested in our attempts to improve encapsulation for the platform developed in this study, yet they did not yield improved outcomes. Ultimately, using the active loading method, the maximum encapsulation efficiency achieved was 21.77 ± 1.8% for the PS liposomes and 18.48 ± 2.34% for the HSPC liposomes (Table S2).

We examined the effect of PEG incorporation on the cellular uptake of liposomes on BxPC-3 and MIA PaCa-2 cell lines using flow cytometry analysis (Figure S4, S5). As expected, in MIA PaCa-2 cells, PEG reduced the gap in the efficacy between HSPC and PS liposomes, when the effect of PEGylated PS liposomes decreased compared to the non-PEGylated PS liposomes. Incorporation of PEG prolongs circulation time of liposomes in the body resulting in improvement in tissue penetration, however at the cellular level, it exhibits significantly lower uptake [36]. Interestingly, in BxPC-3 cells adding PEG to HSPC liposomes revealed a higher uptake than the non-PEGylated HSPC liposomes, while its addition to PS liposomes didn’t lead to any change.

### Uptake of PS Liposomes and The Intracellular Tracking

To examine the uptake kinetics of PS liposomes, we treated MIA PaCa-2 cells with rhodamine labelled PS liposomes over a period of 48 hours (Figure 3). Over that period, the liposomes exhibited increasing cellular accumulation, behaving like an exponential pattern. As long as the liposomes were present in the extracellular space, they tended to enter the cells. Thus, over the examined period, saturation wasn’t observed, which may indicate that the internalization exceeds the degradation and clearance activities. In addition, this pattern of liposomes cellular internalization confirms their stability and implies the PS liposome availability, these properties can certainly contribute to the potential of these particles as drug carriers.

**Figure 3:**
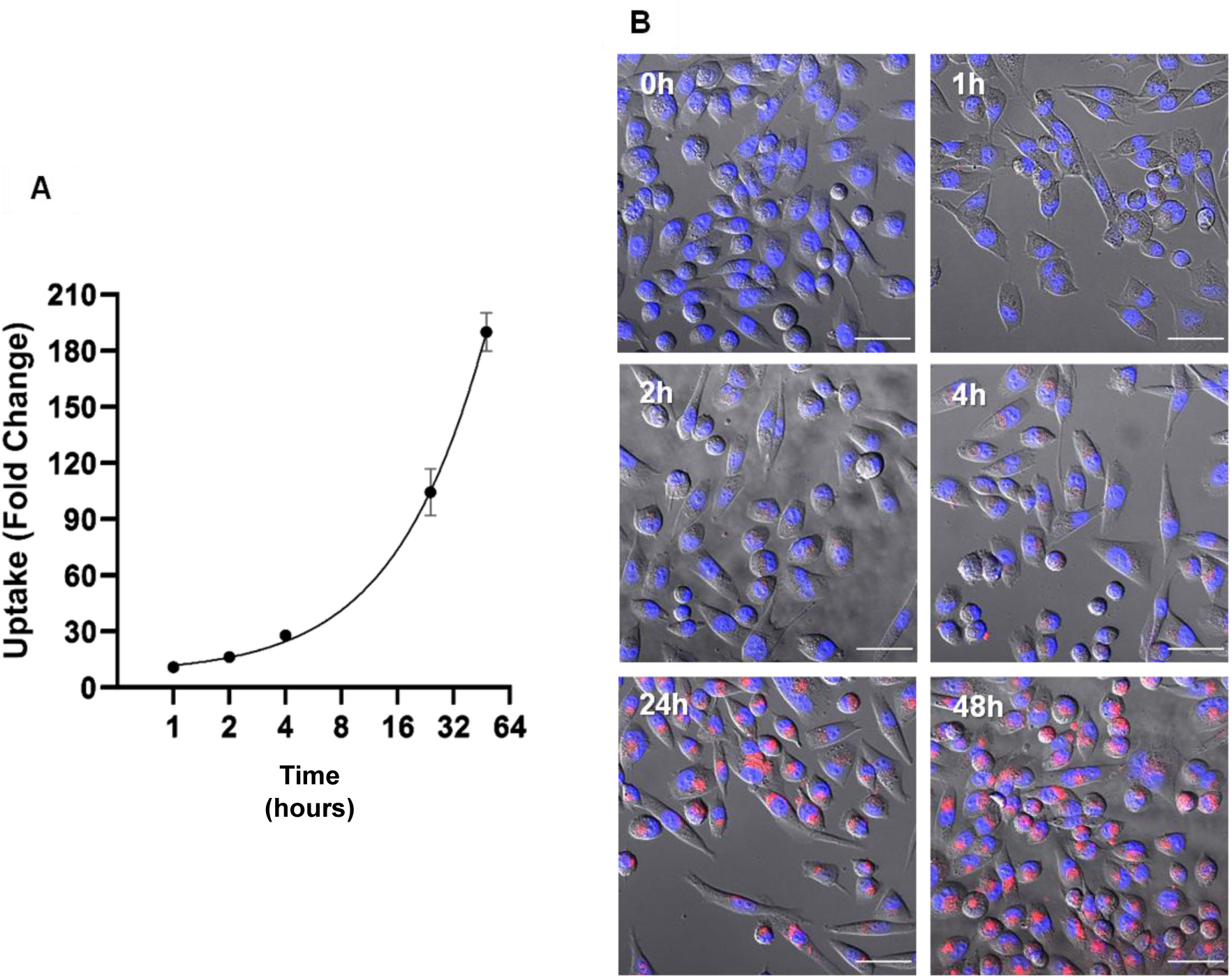
Kinetics of PS liposome uptake by pancreatic cancer cells. **(A)** MIA PaCa-2 cells were treated with rhodamine labelled PS liposomes and the cellular internalization was examined by flow cytometry for the desired time units. The median intensity values were normalized to the unstained values. The solid line between the dots represents the nonlinear regression fitted curve. The dots represent the mean of ± SD (n=3); **(B)** Compatible visualization through confocal images, scale bar 50µm.

Exosomes are internalized by cells through various energy-dependent endocytic mechanisms, such as phagocytosis, micropinocytosis, clathrin-mediated endocytosis, caveolae-dependent endocytosis, and lipid raft-mediated endocytosis [37,38]. It is hypothesized that lipid nanoparticles containing a high concentration of PS are primarily taken up through micropinocytosis [39]. Lowering the temperature to 4°C is known to inhibit energy-dependent uptake processes, including endocytosis [40,41]. Following the methodology of Nagai et al. [41], we investigated the energy-dependent transport of PS liposomes by treating BxPC-3 and MIA PaCa-2 cells with rhodamine labelled PS liposomes at both 4°C and 37 °C (Figure 4). In both cell lines the uptake was completely blocked by reducing the temperature, and the rhodamine intensity matched that of the background, corresponding to the intensity measured in untreated cells. This suggests that the internalization of PS liposomes is driven by active transport mechanisms such as endocytosis. Supporting results were observed for DSPC/Chol/DOPS liposome, when internalization dramatically decreased following incubation at 4 °C [30]. Furthermore, exosome-mimicking liposomes composed of DOPC/SM/Chol/DOPS/DOPE were shown to undergo internalization into human lung cancer cells via caveolae-mediated endocytosis and micropinocytosis [29].

**Figure 4:**
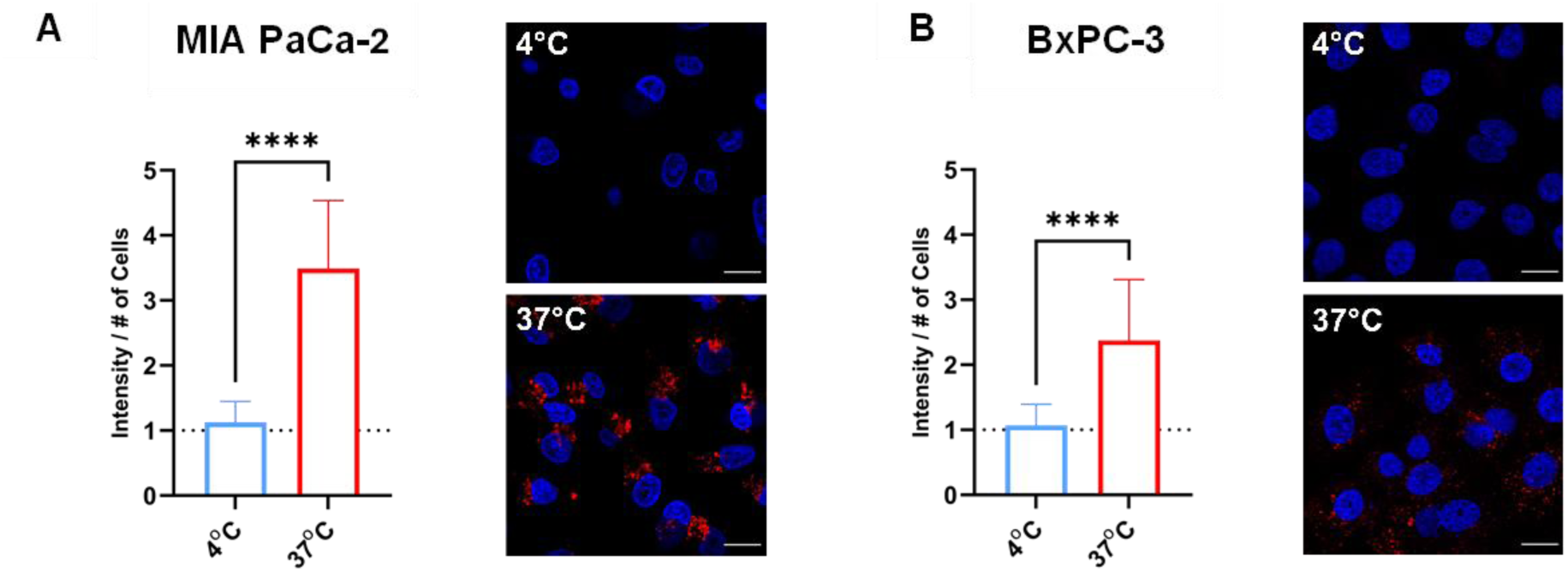
Endocytosis of PS liposome. Confocal microscopy analysis of PS liposome endocytosis into MIA PaCa-2 **(A)** and BxPC-3 **(B)** cells. Cells were incubated with rhodamine-labelled PS liposome for 2 hours in the desired condition. Cellular internalization was evaluated by rhodamine intensity, normalized to cell count, and compared with untreated cells; Scale bar 20 µm. The results are presented as the mean ± SD (n ≥ 13). Statistical analysis was performed using unpaired t-test, significance levels: *\*\*\*\*p* < 0.0001

In addition, we tracked the intracellular route of PS liposomes in MIA PaCa-2 cells by measuring their accumulation in the lysosome and nucleus. Tracking PS liposomes in the lysosome revealed gradual accumulation that began already after 2 hours and peaked at around 24 hours (Figure 5A, Video S1B). These results align with the uptake kinetics, where internalization surpasses cellular clearance. The accumulation of PS liposomes within cells increases, while their colocalization with lysosomes remains limited. This suggests that the intense internalization and accumulation of PS liposomes inside the cell exceed cellular clearance capabilities, highlighting a potential advantageous feature in overcoming drug resistance by effectively increasing the intracellular concentration of the therapeutic payload. The capacity of PS liposomes to mitigate chemoresistance positions these nanocarriers as a promising strategy to enhance the efficacy of chemotherapeutic agents, especially in challenging contexts like pancreatic cancer, where resistance to treatment is common.

**Figure 5:**
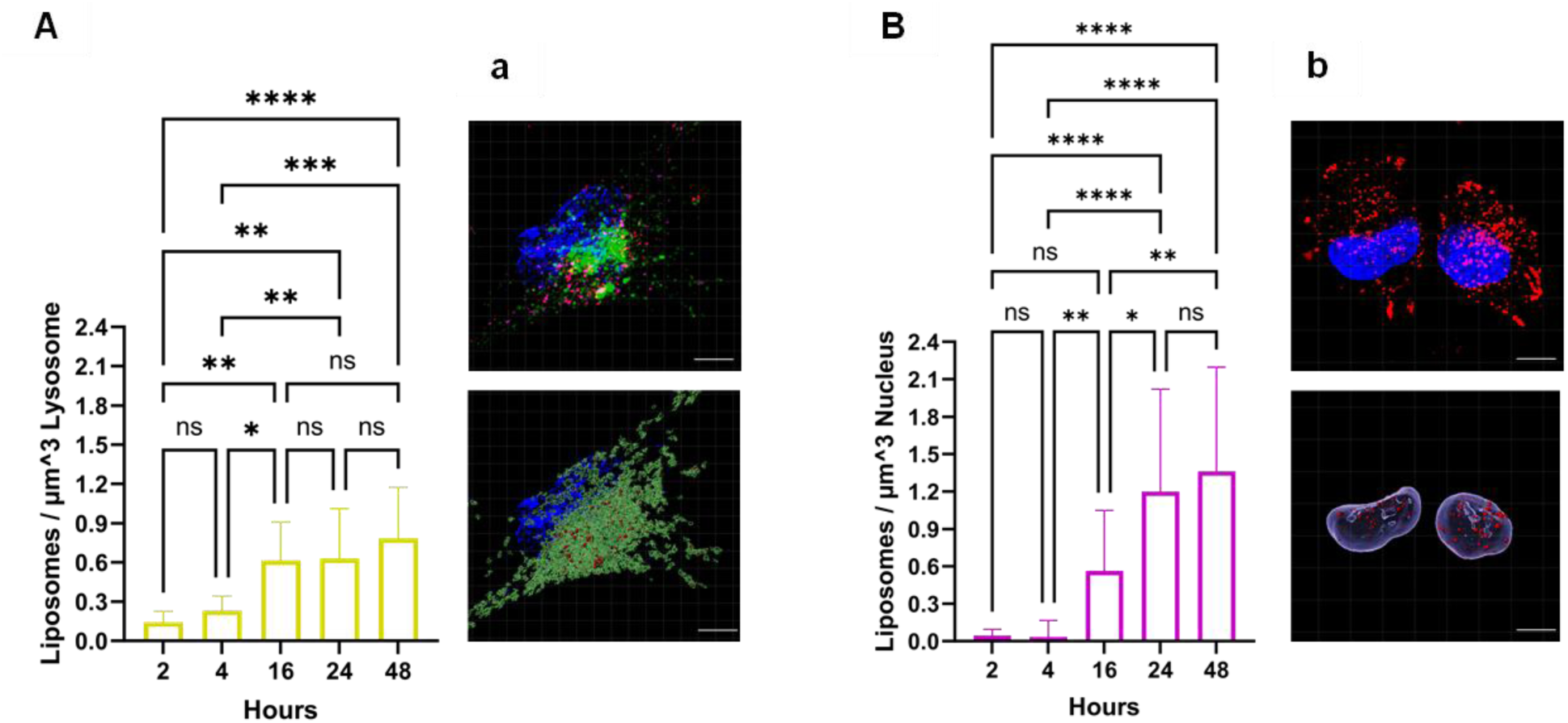
Intracellular tracking of PS liposome. (**A, a)** Colocalization analysis of rhodamine-labelled PS liposomes with lysosomes; **(a)** Representative confocal image captured after 48-hour treatment and its 3D Imaris model, scale bar 10 µm. Liposomes at a distance 0 or less from the lysosome (green) are shown as red dots. Cells were incubated with CellLight™ Lysosomes-GFP, BacMam 2.0 reagent for lysosome staining. Following washing, rhodamine-labelled PS liposomes were introduced for a 2-hour incubation and imaged at each experimental time point. Data is presented as mean ± SD (n ≥ 10), statistical analysis was perfumed using through one-way ANOVA followed by subsequent multiple comparisons test. Significance levels: *\*p* < 0.05, *\*\*p* < 0.01, *\*\*\*p* < 0.001, *\*\*\*\*p* < 0.0001, ns (not significant); **(B, b)** Colocalization analysis of rhodamine-labelled PS liposomes with nucleus; **(b)** Representative confocal image captured after 24-hour treatment and its 3D Imaris model, scale bar 10 µm. Liposomes at a distance 0 or less from the nucleus (blue) are shown as red dots. Cells were incubated with rhodamine-labelled PS liposomes for the compatible experimental time point; For comparison, the ratio of points within zero or less distance from the lysosome or nucleus to the compatible organelle’s volume was calculated. The results are displayed as mean ± SD (n ≥ 14), with statistical analysis conducted through Kruskal-Wallis test followed by multiple comparisons test. Significance levels: *\*p* < 0.05, *\*\*p* < 0.01, *\*\*\*\*p* < 0.0001, ns (not significant).

While attempting to monitor PS liposomes routes within the cells, we discovered that the liposomes demonstrated an intriguing tendency for attaching themselves to the nucleus surface (Figure 5B, Video S1A). The liposomal accumulation on the nucleus was neglectable in the first few hours but then the colocalization started to increase over time, within the examined time frame. This nuclear affinity can be explained by the interaction between PS and nucleosome histones as PS colocalize with epichromatin in interphase nuclei and mitotic chromosomes [42]. This colocalization underscores PS’s critical role in the architecture of cellular nuclei, notably in the reformation of the nuclear envelope and the organization of chromatin during the cell cycle. Separately, evidence exists for the natural exosomes’ ability to target the nucleus. For example, exosomes derived from M2 macrophages exhibited internalization into the nucleus [10]. In an additional study, exosomes from kidney cells demonstrated the capability to reach the nucleus, with observed entry increasing over time [43]. In our platform, the potential to deliver gemcitabine to the nucleus may underscore a critical aspect of its intracellular mechanism of action. Gemcitabine, also called 2’,2’-difluoro2’-deoxycytidine or dFdC, is transported into cells predominantly through human nucleoside transporters (hNTs), specifically the equilibrative (ENTs) and concentrative (CNTs) varieties [44,45]. Within the cytoplasm, gemcitabine is phosphorylated in three steps to reach its active metabolite form. Initially, deoxycytidine kinase (dCK) phosphorylates it to form dFdCMP. This is followed by its conversion to gemcitabine diphosphate (dFdCDP) through phosphorylation by nucleotide monophosphate kinase (NMPK). Finally, the nucleotide diphosphate kinase (NDPK) produces the triphosphate product, dFdCTP [45–47]. dFdCTP disrupts DNA synthesis, ultimately resulting in cell death [46,48]. Theoretically, the delivery of gemcitabine into the nucleus via liposomes could permit its action at the nuclear level. Concurrently, the accumulation of liposomes in the cytosol allows for the activity of gemcitabine in this compartment as well. This dual localization strategy could potentially potentiate the drug’s effectiveness by engaging multiple intracellular targets.

### Dose Response

To assess the pharmacology efficacy of gemcitabine loaded PS liposomes, we compared the effect on cellular viability to the basic formulation and to free gemcitabine. We tested five gradual concentrations of gemcitabine in order to produce IC50 graphs for each one of the treatments. Despite the slight advantage of HSPC in the maximum effect in both cell lines (Figure 6), the PS liposomes exhibited more potency to the cells, which is presented by sigmoidal IC50 graphs. For the MIA PaCa-2 cell line, PS liposomes showed IC50 values of 14.19 µM after 24 hours of treatment and significantly decreased to 1.357 µM following 48 hours of exposure (Figure 6A-B). In contrast, HSPC liposomes had IC50 values of 129.5 µM at 24 hours, which reduced to 17.27 µM after 48 hours (Figure 6A-B). Similarly, in BxPC-3 cells, PS liposomes were more potent, with IC50 values of 1.494 µM at 24 hours, decreasing sharply to 0.1977 µM at 48 hours (Figure 6C-D). HSPC liposomes presented higher IC50 values of 9.909 µM at 24 hours and 0.9582 µM at 48 hours (Figure 6C-D). These findings suggest that PS liposomes are more effective at lower concentrations and over time.

**Figure 6:**
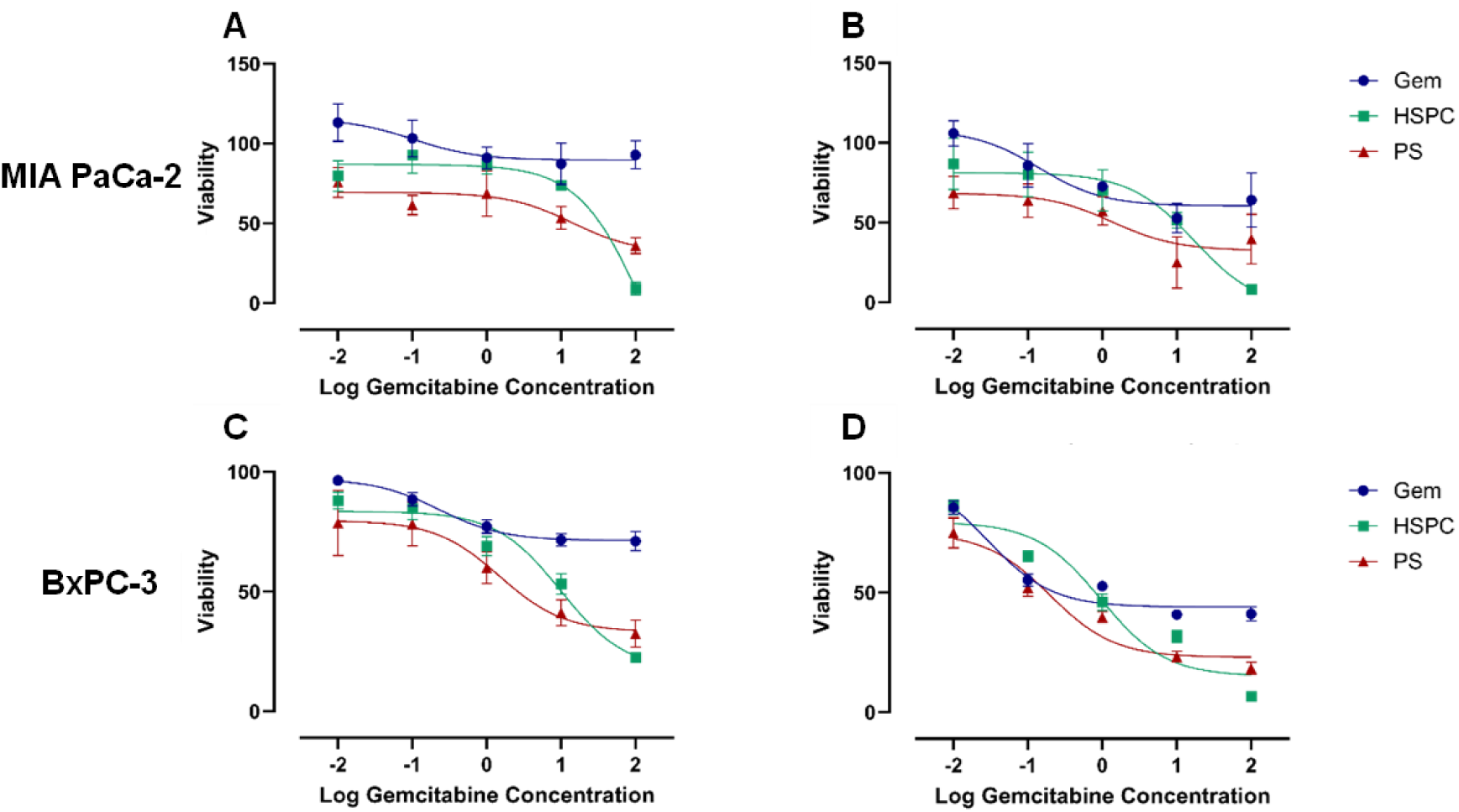
Dose Response. MIA PaCa-2 **(A, B)** and BxPC-3 **(C, D)** cells were treated with gemcitabine encapsulated liposomes and free drug to produce dose response curves. The cells were treated for 24-**(A, C)** and 48-hours **(B, D)** and cell viability was examined by XTT. Nonlinear regression was analyzed for each treatment to produce inhibition dose response curves. The results are presented as the mean of ± SD (n=5).

### Programmed Cell Death Caused by PS Liposomes

To investigate whether the drug delivery of liposomes induces programmed cell death, we assessed cellular expression of cleaved PARP1 using Western Blot analysis. For this purpose, cells were exposed to free gemcitabine and gemcitabine encapsulated PS and HSPC liposomes, at identical drug concentrations for durations of 24- and 48-hours. For MIA PaCa-2 the effect of PS liposomes, like the free drug, resulted in high expression of cleaved PARP1 as compared to the untreated cells (Figure S6A). This tendency was amplified in the 48-hour treatment (Figure 7A). However, the treatment of HSPC liposomes didn’t promote any expression and the cleaved PARP1 level remained as in the control group. In contrast, the results of BxPC-3 didn’t show any difference, as all the treatments similarly upregulated the expression of cleaved PARP1 (Figure S6B).

**Figure 7:**
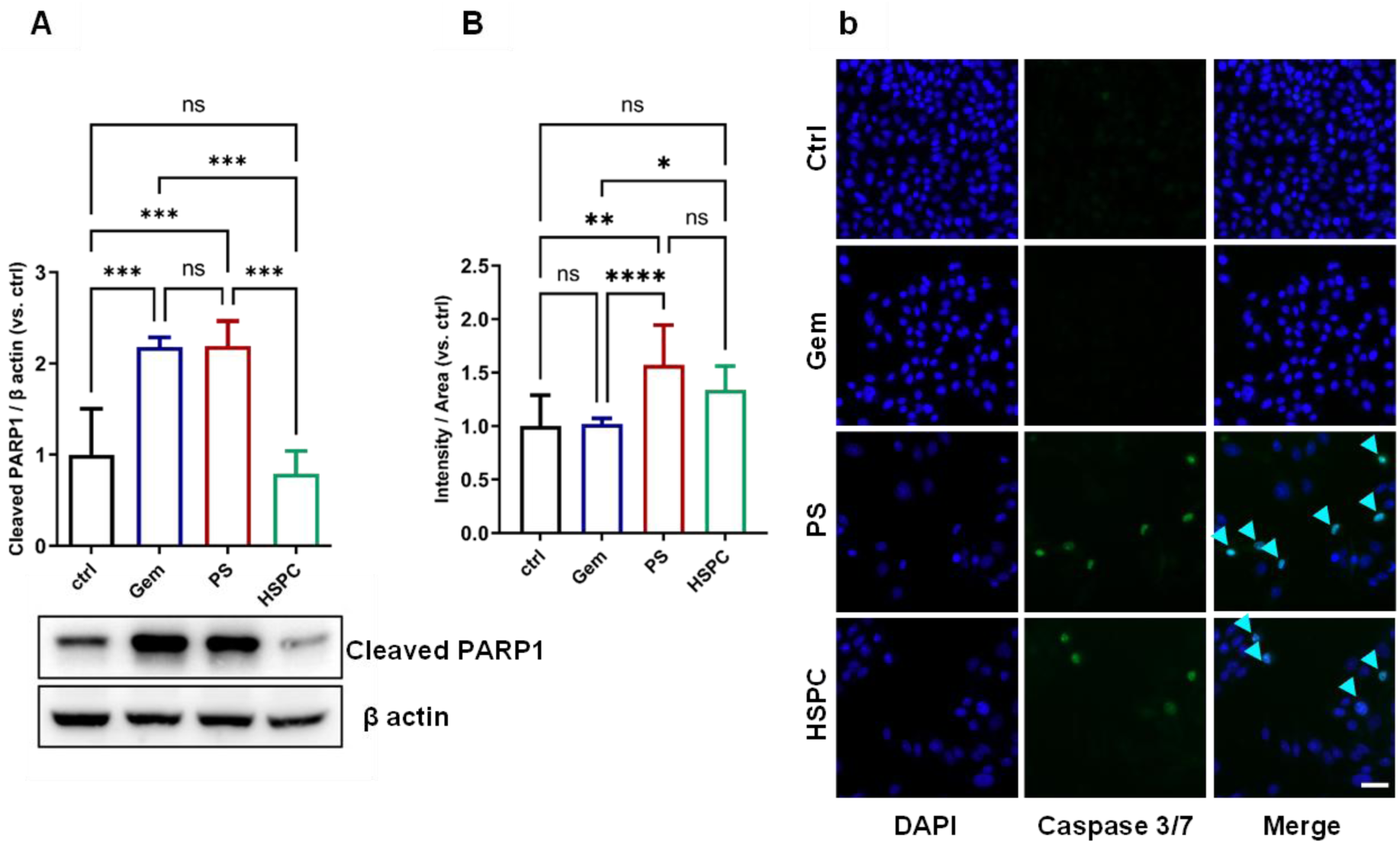
Induction of Programmed Cell Death in Pancreatic Cancer Cells by PS Liposomes. (A) Western blot analysis of cleaved PARP1 expression after 48-hour treatment with gemcitabine-encapsulated liposomes and free drug in Mia Paca-2 cells. The results are presented as the mean ± SD (n ≥ 3). Statistical analysis was performed using one-way ANOVA followed by subsequent multiple comparisons test. Significance levels: *\*\*\*p* < 0.001, ns (not significant); (B) Caspase 3/7 activity was assessed in Mia Paca-2 cells using CellEvent detection reagent under the same experimental conditions as the cleaved PARP1 analysis. The detection reagent was introduced, and cells were visualized using fluorescence microscopy (b), Scale bar 50 µm. Results are displayed as the mean ± SD (n > 12), and statistical analysis was conducted through one-way ANOVA followed by subsequent multiple comparisons test. Significance levels: *\*p* < 0.05, *\*\*p* < 0.01, *\*\*\*\*p* < 0.0001, ns (not significant). conducted through one-way ANOVA followed by subsequent multiple comparisons test. Significance levels: *\*p* < 0.05, *\*\*p* < 0.01, *\*\*\*\*p* < 0.0001, ns (not significant).

Next, we investigated Caspase 3/7 as an additional marker for programmed cell death, given that both caspase-3 and caspase-7, cysteine proteases, are activated during apoptosis and initiate the final execution pathway [49,50]. In the case of BxPC-3 (Figure S7B), an elevation in caspase 3/7 levels was observed following treatment with PS liposomes, and this elevation showed a slight enhancement after 48 hours of treatment. While a similar trend was evident for HSPC liposomes, the expression levels under the 48-hour treatment of the PS liposomes exhibited statistical significance when compared to both free gemcitabine and the untreated control cells. In the MIA PaCa-2 cells, a 24-hour treatment of PS liposomes demonstrated a notable increase in caspase 3/7 levels compared to the other treatments, which did not exhibit any elevation (Figure S7A). This increase further intensified after 48 hours, while HSPC liposomes also led to an upregulation of expression (Figure 7B).

### Tumor Growth Inhibition by PS Liposomes

To investigate the treatment efficacy of PS liposomes in pancreatic cancer, we administered Gemcitabine-PS liposomes at a concentration of 10 mg/kg body weight. Due to the limitation of the encapsulation capacity of the liposomes, we selected the maximum feasible dose of gemcitabine for the delivery. We evaluated its effect on tumor growth by comparing it to a 10-fold higher concentration of free gemcitabine, as well as to a free drug at the same encapsulated dose concentration. Throughout the treatment with PS-liposomes, no indications of toxicity were observed, and the mice’s weight exhibited a gradual increase, as in the control and free drug groups (Figure S8). Gemcitabine-PS liposomes demonstrated effective inhibition of tumor growth (p<0.05), what was not observed in the free drug groups (Figure 8B, C). Although the inhibition effect of the liposomes exhibited superiority over the 100 mg/kg free gemcitabine treatment, a statistical analysis between the two groups did not reveal a significant difference. (*p*=0.0658) (Figure 8C).

**Figure 8:**
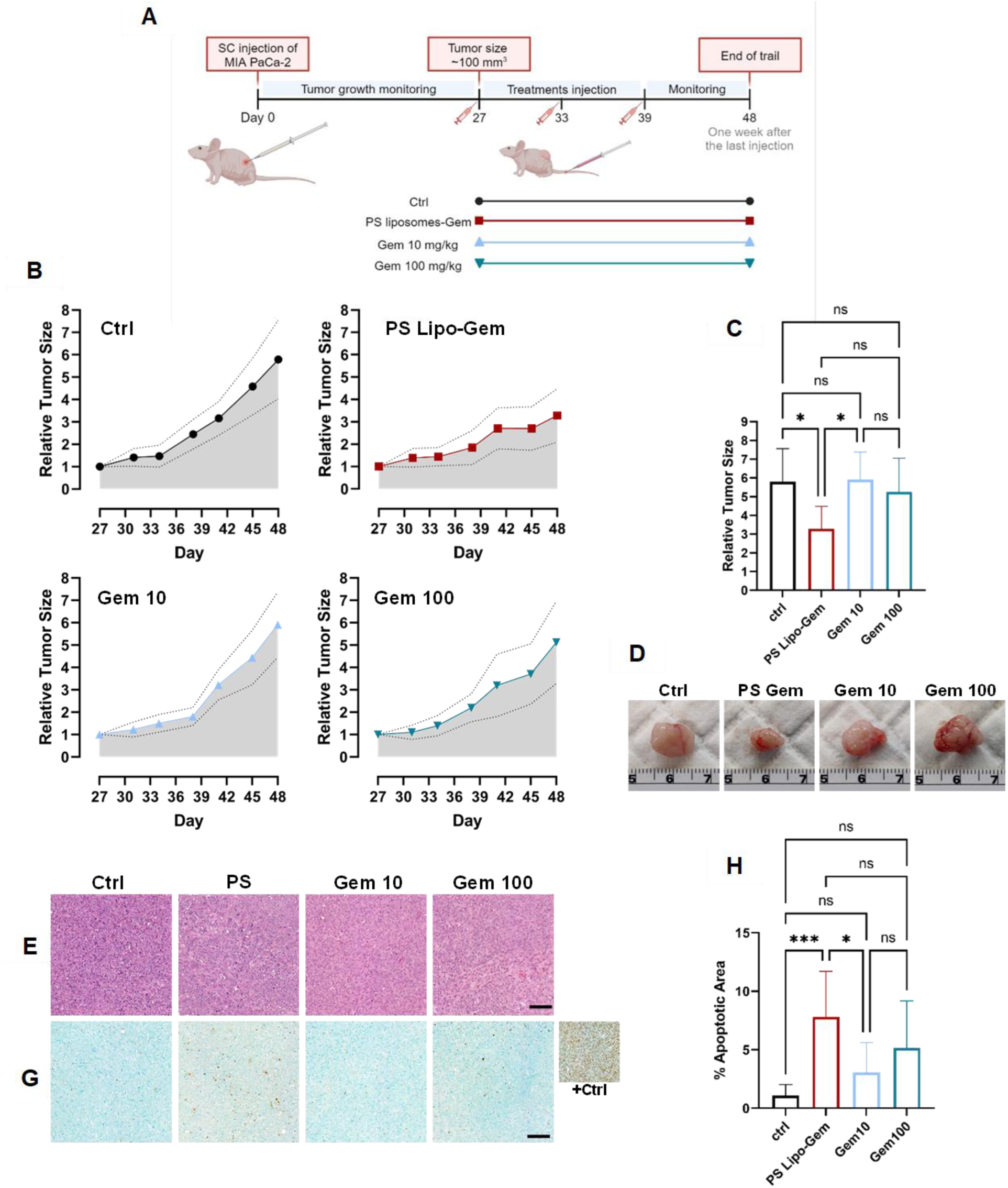
Tumor Growth Inhibition by PS Liposomes. **(A)** Scheme of *in vivo* study design; (B)Tumor-bearing mice received intravenous treatment with gemcitabine-loaded PS liposomes (10mg/kg gemcitabine), free gemcitabine (10 and 100 mg/kg), and a 5% D-glucose solution control. Tumor sizes were assessed every 2-3 days and normalized to the initial day’s size. Graphs present mean ± SD (4≤n≤6); **(C)** Gemcitabine-loaded PS liposomes significantly inhibited tumor growth compared to the control and free gemcitabine. Data is presented as the mean ± SD. Statistical analysis was conducted using one-way ANOVA. Significance level: *\*p* < 0.05. **(D)** Tumors were harvested at the experiment’s end; **(E)** Histology analysis performed, scale bar: 200 µm.; **(G)** TUNEL staining, scale bar: 200 µm; **(H)** Gemcitabine-loaded PS liposomes increased the apoptosis expression in tumor sections compared to control and free gemcitabine. Mean ± SD, statistical analysis via one-way ANOVA followed by subsequent multiple comparisons test. Significance level: *\*p* < 0.05, *\*\*\*p* < 0.001, ns (not significant).

About seven weeks after tumor induction, mice were sacrificed, and the tumors were resected for performing H&E and TUNEL staining. Swelling cells and massive necrosis were observed in the tumor tissues from Gemcitabine-PS liposome treated mice, as revealed by H&E staining (Figure 8E). In contrast, there were not significant changes in tumors of free drug treated mice, compared to the control. TUNEL staining revealed a significantly elevated level of apoptotic expression in tumor sections following treatment with PS liposomes (Figure 8G-H). This increase in apoptosis was markedly higher than that in the control group and exceeded the levels observed in the group treated with an equivalent concentration of free gemcitabine. Furthermore, the apoptotic activity induced by PS liposomes was greater to a certain extent compared to the group treated with ten times consternated free gemcitabine that encapsulated within the liposomes (Figure 7H).

### Biodistribution of PS Liposomes

Next, we aimed to monitor the biodistribution of PS liposomes and to evaluate their tumor accumulation efficacy. To this end, PS liposomes were compared with HSPC liposomes (Figure 9). We conducted the examination using rhodamine labelled liposome for fluorescence detection using IVIS. Consistent with our expectations, both types of liposomes demonstrated high level of accumulation in the liver, with statistically insignificant preference for PS liposomes (mean radiant efficiency value of PS liposomes 9.65e+07 (p/s/cm²/sr) / (µW/cm²) ± 7.47e+07 and for HSPC liposomes 3.73e+07 (p/s/cm²/sr) / (µW/cm²) ± 1.69e+07; p=0.1226). Notably, meaningful tumor accumulation was observed in both treatments, showing no significant differences between them. However, in all other examined tissues, a higher level of HSPC liposome accumulation was noted in comparison to PS liposomes.

**Figure 9:**
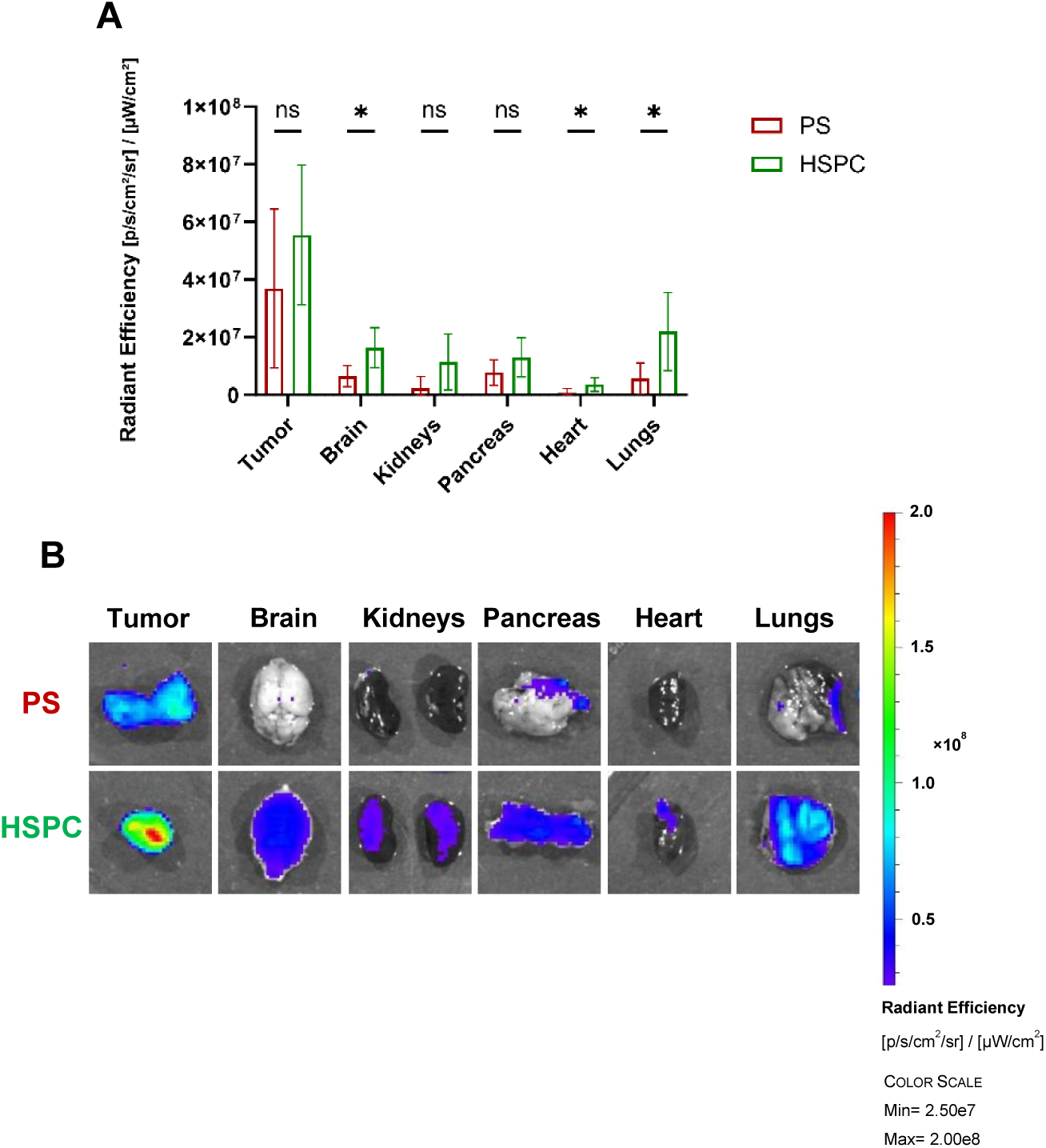
Biodistribution of PS liposomes. **(A)** *in vivo* biodistribution comparison of PS and HSPC rhodamine-labelled liposomes. Mice received intravenous injections of the liposomes and were euthanized four hours post-injection for organ analysis using IVIS imaging. The results are presented as the mean ± SD (n=6). Statistical analysis was performed using unpaired t-test, significance levels: *\*p* < 0.05, ns (not significant); **(B)** Representative fluorescence imaging of the tissues.

This finding suggests that PS liposomes may exhibit greater tumor specificity, whereas HSPC liposomes appear to have a more random distribution. Despite the slight accumulation of PS liposomes in non tumor organs such as the brain, pancreas, and lungs, histopathological evaluations have confirmed that the delivery of gemcitabine via PS liposomes does not cause any damage to these organs (Figure 10C). This lack of detrimental effects supports the potential of PS liposomes for safe drug delivery, emphasizing their capability for targeted therapy with minimal off-target risks. Further research could explore optimizing their composition and dosage to enhance tumor specificity even further.

**Figure 10:**
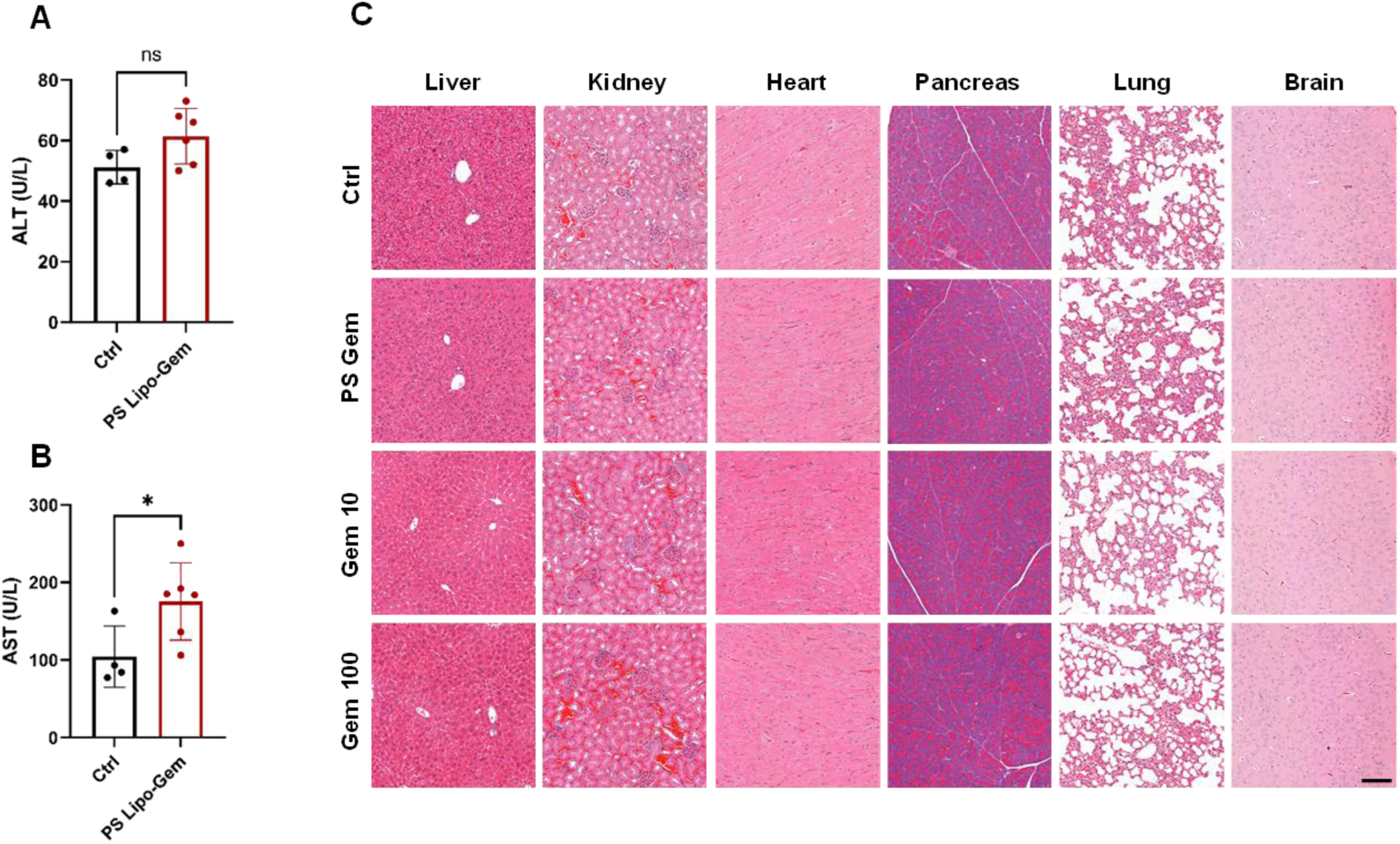
Assessment of Liver Function and Toxicity Following PS Liposome Treatment. (A) Alanine aminotransferase (ALT) and (B) Aspartate aminotransferase (AST) blood levels, were assessed post PS liposome treatment and compared to the control group. Data is shown as mean ± SD (4≤n≤6). Statistical analysis was performed using unpaired t-test. Significance level: *\*p* < 0.05, ns (not significant); (C) Histopathological evaluation of various organs was conducted using hematoxylin and eosin (H&E) staining, scale bar 200 µm.

### Examination of Liver Function and Toxicity

Liposome accumulation in the liver is a well-documented phenomenon, with the liver acting as a major clearance barrier for substances present in the systemic blood circulation [51]. As mentioned above, the biodistribution examination revealed a high accumulation of PS liposomes in the liver. This observed tendency prompted further investigation into the treatment’s effect on liver function. For this purpose, we collected blood serum samples from the control and gemcitabine-PS liposomes groups to diagnose liver function. In Gemcitabine-PS liposomes mice, both ALT and AST levels exhibited a slight increase compared to the control mice (Figure 10A-B). The rise in ALT levels was relatively milder compared to AST levels. Importantly, ALT and AST levels in both treatment groups stayed within the normal range (ALT 27-78 U/L and AST 50-215 U/L), according to established norms of nude mice biochemistry from © Charles River Laboratories. Indeed, the histopathological analysis through H&E staining did not reveal any notable histological distinctions among the groups (Figure 10C).

In addition to the liver tissues, H&E staining was performed for examination of systemic toxicity. The brain, heart, kidney, lungs, and pancreas were excised after the mice were euthanized and their sections were stained. Any distinguishable abnormalities were observed, indicating negligible systemic toxicity (Figure 10C).

## Conclusions

In this study, we established a novel and fundamental therapeutic platform for pancreatic cancer treatment, focusing on the application of exosomal lipid composition mimicry. Driven by the discovery that exosomes derived from M2 macrophages are preferentially internalized by cancer cells within the pancreatic tumor microenvironment, we concentrated on analyzing the unique lipid components of the exosome membrane. Interestingly, among all phospholipids identified, incorporating PS into a basic liposome formulation yielded the highest cellular uptake. Our developed synthetic exosomes, termed PS liposomes, demonstrate superior stability, efficacy, and preferential affinity for pancreatic cancer cells compared to the basic liposome formulation. Significantly, they have been shown to trigger apoptosis in pancreatic cancer cells both *in vitro* and *in vivo*. Unlike the basic liposome formulation, which exhibited a more dispersed biodistribution, PS liposomes displayed enhanced tumor specificity, leading to reduced off-target accumulation in non-tumorous organs. This targeted delivery capability minimizes potential systemic toxicity and side effects, marking an advance in cancer therapy. Moreover, our study indicates that PS liposomes surpass free gemcitabine in inhibiting tumor growth.

This achievement represents a significant leap forward in drug delivery for pancreatic cancer, a field desperately in need of innovative therapeutic strategies. Despite existing treatments, pancreatic cancer patients’ survival rates remain low. The development of synthetic exosomes as a highly effective drug delivery platform introduces a promising avenue for improving patient outcomes, warranting further investigation into its underlying mechanisms and future applications.

Exploring the lipid composition of exosomes, a relatively underexplored area compared to their protein and nucleic acid contents, has illuminated the potential of these components in enhancing synthetic drug delivery systems’ functionality. Further research is essential to refine and expand the capabilities of these synthetic exosomes. The need for further optimization, modifications to enhance delivery efficiency, and testing the encapsulation of different chemotherapeutic agents for synergistic effects is clear. Additional studies should explore the applicability of PS liposomes in other cancer types and investigate the long-term effects and safety in clinical settings. Expanding exosomal mimicry through multidisciplinary studies, particularly by incorporating targeting ligands or surface proteins for improved specificity to pancreatic cancer and tissue penetration, could be highly beneficial. The success of FDA and EMA-approved liposomal products, along with the established safety of phosphatidylserine as a dietary supplement, lays a solid foundation for our findings’ clinical application.

In conclusion, our work highlights the significant potential of leveraging exosomal lipid profiles for cancer therapy. By introducing an innovative drug delivery system that mimics the natural properties of exosomes, we propose a novel and effective approach to treating pancreatic cancer, setting the stage for a future where this devastating disease can be addressed more effectively.

## Supporting Information

- Supporting information includes graphical presentation of lipidomic analysis findings, liposomes characterization, description of gemcitabine encapsulation, cellular uptake of liposomes, apoptosis expression and additions to in vivo experiments (PDF).
- Video of nuclear colocalization produced by IMARIS software (mp4).
- Video of lysosome colocalization produced by IMARIS software (mp4).

## Author Contributions

**TNS** conceived and designed the study, performed the experiments, analyzed the data, prepared the figures, and wrote the manuscript. **AS** provided professional support and scientific guidance, contributed to the study design, and assisted with resources and manuscript review. **LG** conducted the initial lipidomic experiments and examining the uptake of different lipids. **MA** contributed to the execution and analysis of the animal experiments. **NRR** and **OA** contributed valuable insights and manuscript review. **HW** contributed conceptual guidance and mentorship, provided key experimental resources, and critical manuscript feedback. **ZG** provided overall supervision, conceptual guidance, and critical review. All authors reviewed and approved the final manuscript.

## Acknowledgments

Graphical images were made by BioRender.com.

