## Supporting Information for "Synthetic Mimetics of Exosomal Lipid Profile Enhance Gemcitabine Delivery in Pancreatic Cancer"

**Table S1:** Characterization of the different liposome formulations by Size, polydispersity index (PDI) and zeta potential.

| Formulation | Size [nm] | PDI | Zeta potential [mV] |
|---|---|---|---|
| **HSPC** | 100.2 ±1.32 | 0.043 ±0.025 | -9.14 ± 0.70 |
| **PE** | 98.84 ±1.38 | 0.059 ±0.016 | -10 ±1.07 |
| **PI** | 94.41 ±0.51 | 0.048 ±0.018 | -42.7 ±1.28 |
| **PS 5%** | 99.73 ±0.47 | 0.05 ±0.011 | -47 ±0.71 |
| **PS 10%** | 99.78 ± 1.68 | 0.042 ±0.014 | -49.1 ±1.32 |
| **SM** | 89.7 ±0.71 | 0.061 ±0.004 | -9.04 ±4.03 |
| **LPC** | 97.08 ±1.22 | 0.053 ±0.015 | -9.8 ±0.29 |
| **Cer** | 96.6 ±0.22 | 0.042 ±0.011 | -9.03 ±0.95 |

**Table S2:** Characterization of PEGylated HSPC and PS liposome by size, polydispersity index (PDI), particle concentration and zeta potential. The size was verified by measuring the particle diameters in cryo-TEM images. The gemcitabine (Gem) encapsulation efficiency was calculated as the ratio of the measured drug concentration in the liposomes by HPLC to the total loaded drug concentration.

| Formulation | Size [nm] | SD | PDI | Concentration [particles/ml] | Zeta Potential [mV] | SD | Size [nm] - Cryo TEM | SD | Gem encapsulation efficiency % | SD |
|---|---|---|---|---|---|---|---|---|---|---|
| **HSPC** | 96.69 | 2.03 | 0.056 | 1.36E+13 | -40.89 | 2.98 | 94.20 | 2.25 | 18.87 | 2.15 |
| **PS** | 98.04 | 2.65 | 0.052 | 1.36E+13 | -52.42 | 7.37 | 96.13 | 1.67 | 20.35 | 2.20 |

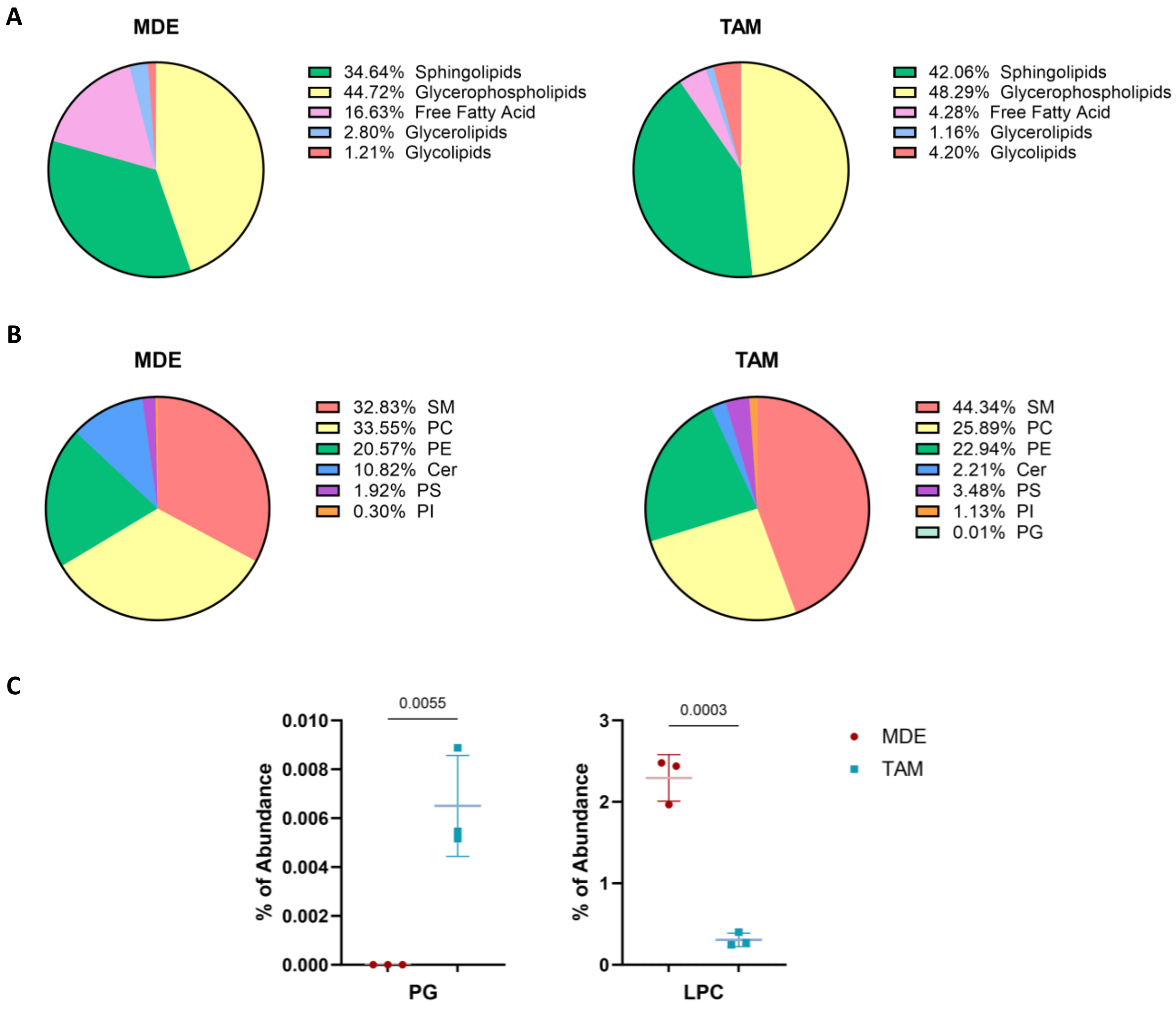


**Figure S1: Differential lipid composition in exosomes derived from M2 polarized macrophages compared to origin cells.** Nearly 210 lipid species from diverse lipid families, including phospholipids, glycolipids, glycerolipids, and free fatty acids, were identified through liquid chromatography-mass spectrometry analysis. **(A)** The total lipid composition in macrophages derived exosomes (MDE) and tumor associated macrophages (TAM) are presented by pie charts. **(B)** Phospholipids distribution in MDE and TAM. Sphingomyelin (SM), phosphatidylcholine (PC), phosphatidylethanolamine (PE), ceramide (Cer), phosphatidylserine (PS) and phosphatidylinositol (PI). **(C)** Comparison of the phosphatidylglycerol (PG) and lysophosphatidylcholine (LPC) relative abundance in MDE and TAM . Three independent biological samples of MDEs and TAMs are presented as mean ± SD. For the comparison of each phospholipid in both samples, unpaired t-test was used as statistical analysis. Significance levels (*p* value) are shown in the graphs.

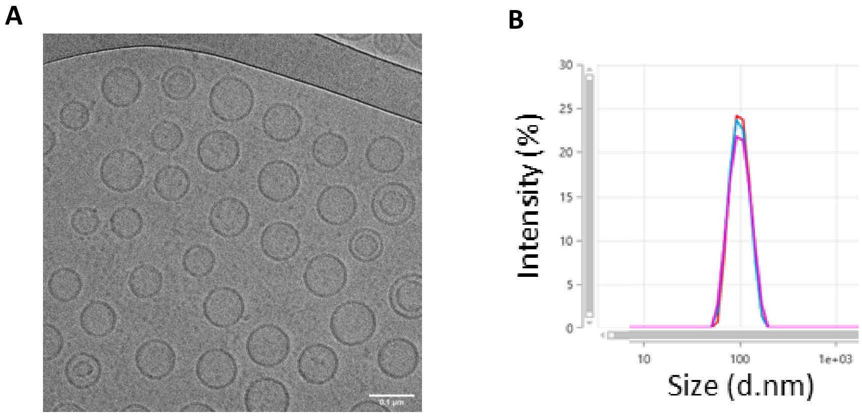


**Figure S2: Characterization of PEGylated HSPC liposome. (A)** Representative cryo-TEM image of HSPC liposomes, scale bar, 0.1 μm; **(B)** HSPC liposomes' size was measured by dynamic light scattering.

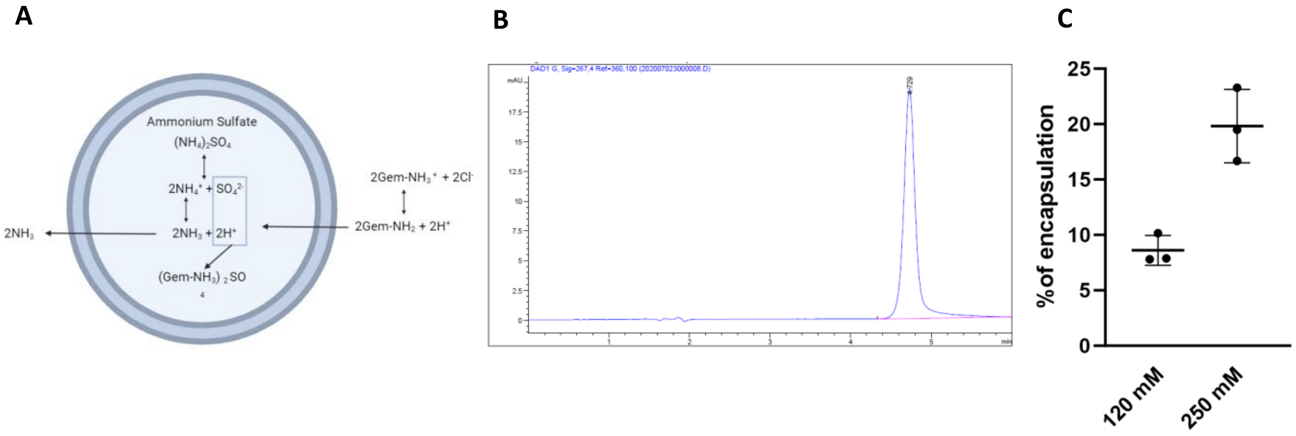


**Figure S3: Gemcitabine encapsulation in PEGylated PS liposome. (A)** 100 nm PS liposomes were loaded with gemcitabine using ammonium sulfate gradient (created in BioReander.com); **(B)** Representative HPLC chromatogram depicting gemcitabine encapsulated within PS liposomes; **(C)** Gemcitabine encapsulation efficiency showed improvement when using a 250 mM ammonium sulfate gradient compared to a 120 mM ammonium sulfate gradient. The data is presented as the mean ± SD (n=3).

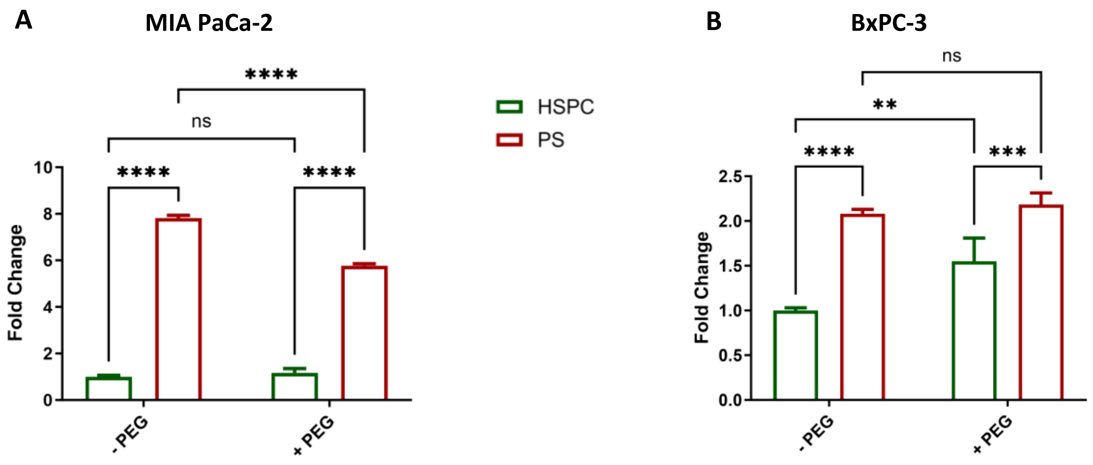


**Figure S4: Uptake of PEGylated and unPEGylated liposomes by pancreatic cancer cells. (A)** Mia PaCa-2 **(A)** and BxPC-3 **(B)** cells were treated with PEGylated (+ PEG) and unPEGylated (- PEG) rhodamine-labelled liposomes for two hours at identical particle concentration. Comparison of the cellular uptake was assessed using flow cytometry. The median intensity values were normalized to those of unPEGylated HSPC. The results are presented as the mean of ± SD (n=3). Statistical analysis was conducted using a two-way analysis of variance (ANOVA) with a subsequent multiple comparisons test; significance levels: $^{**}p < 0.01$, $^{***}p < 0.001$, $^{****}p < 0.0001$, and ns (not significant).

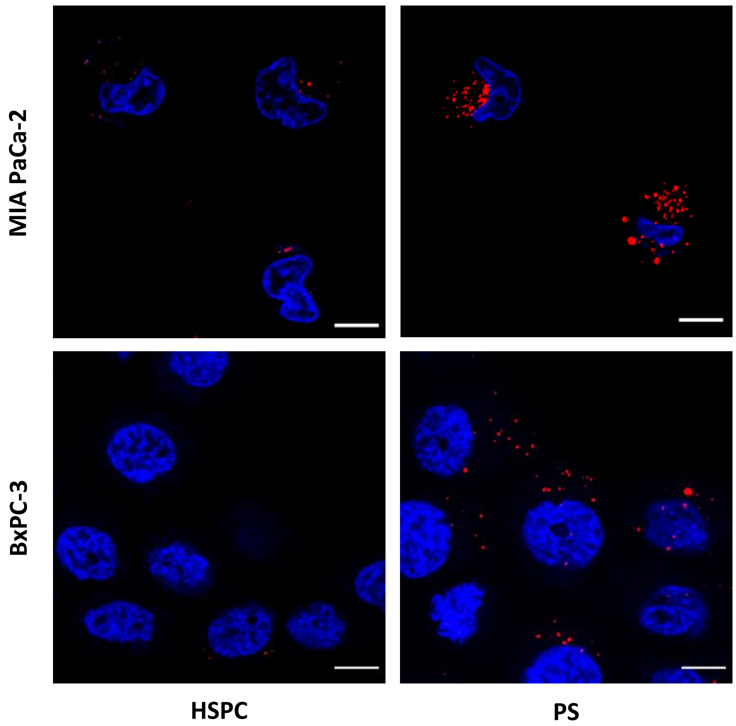


**Figure S5: Internalization of PEGylated liposomes into pancreatic cancer cells.** The intracellular internalization of the PEGylated liposomes was observed by confocal microscopy. Mia PaCa-2 and BxPC-3 cells were treated with PEGylated rhodamine-labelled liposomes for two hours at identical particle concentration, scale bar 10μm.

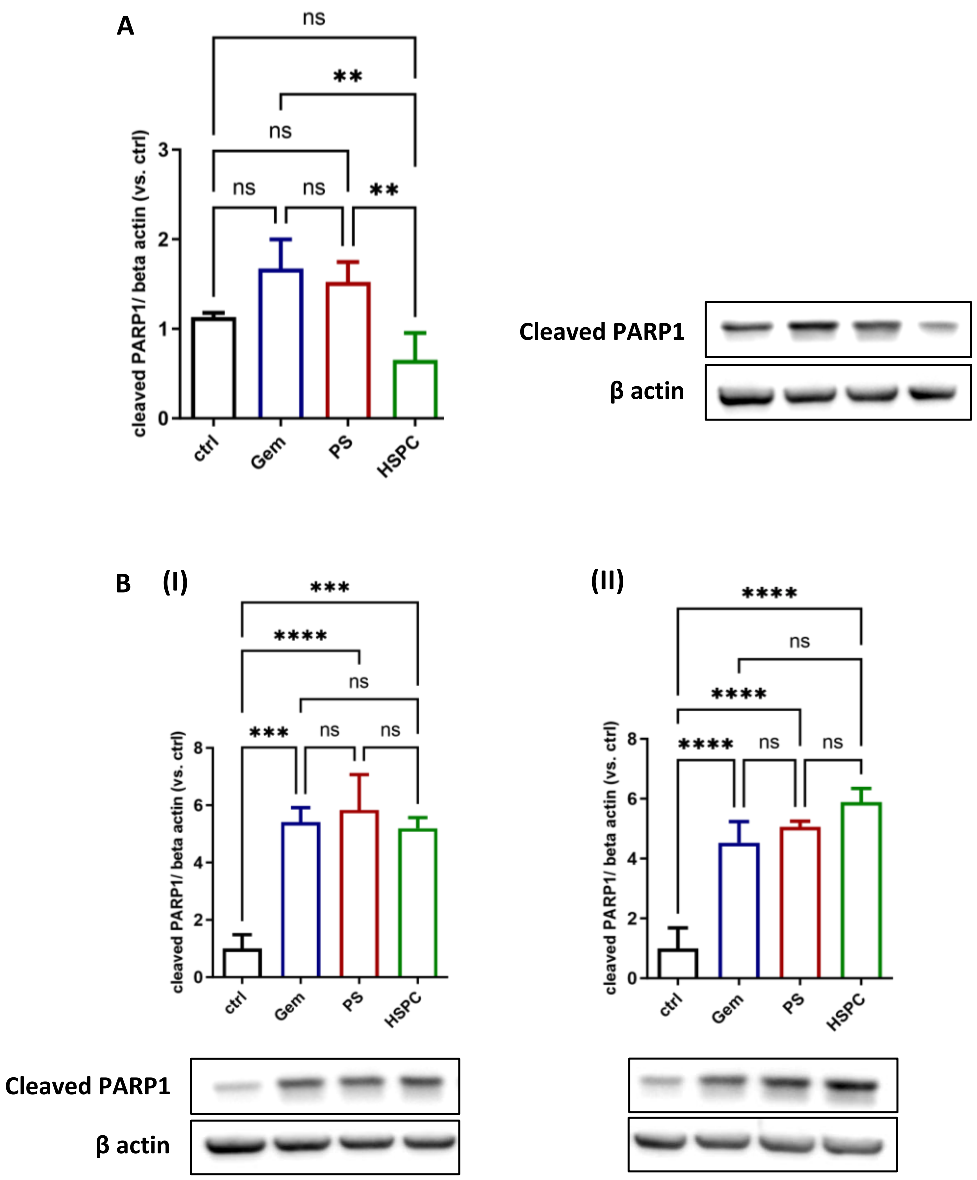


**Figure S6: Cleaved PARP1 expression in pancreatic cancer cells.** Western blot analysis of cleaved PARP1 expression in response to gemcitabine-encapsulated liposomes and free drug. **(A)** 24- hour treatment in MIA PaCa-2 cells; **(B)** cleaved PARP1 expression in BxPC-2 cells after 24- (I) and 48- hour (II). Data is represented as mean ± SD (n ≥ 3). Statistical analysis was conducted using one-way ANOVA followed by subsequent multiple comparisons test. Significance levels: ****$p < 0.0001$, ***$p < 0.001$, **$p < 0.01$, ns (not significant).

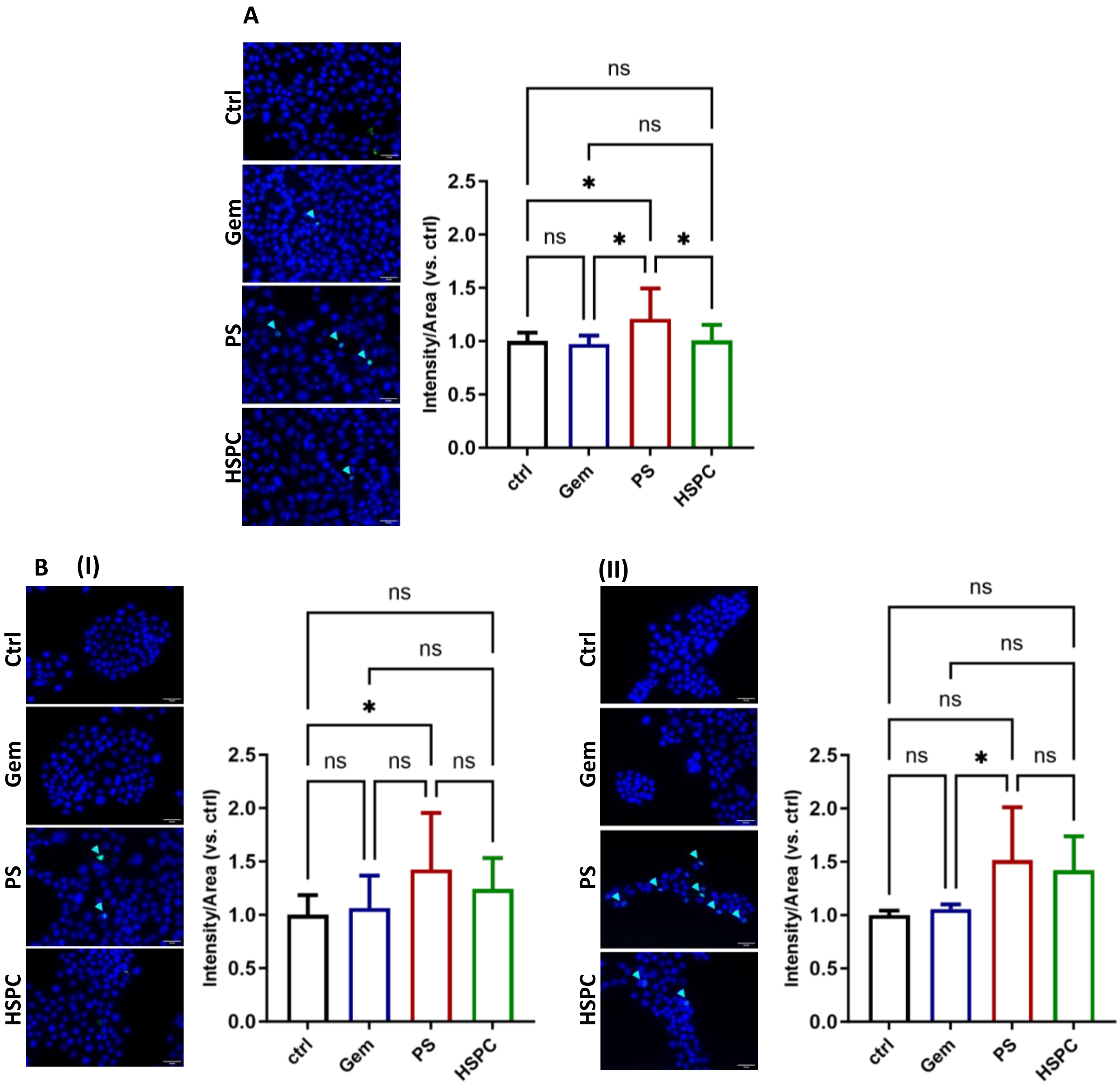


**Figure S7: Caspase 3/7 expression in pancreatic cancer cells.** Utilizing fluorescence microscopy, caspase 3/7 expression was examined in response to gemcitabine-encapsulated liposomes and free drug treatment. Following 24- and 48-hour treatments, MIA PaCa-2 and BxPC-3 cells were exposed to a caspase 3/7 detection reagent, fixed, and subjected to imaging. **(A)** 24- hour treatment in MIA PaCa-2; **(B)** BxPC-3 cells after 24- (I) and 48-hour (II) treatments. The intensity of caspase 3/7 within nucleus areas was quantified. Data is presented as mean ± SD ($n \geq 7$). Statistical analysis involved a one-way ANOVA followed by subsequent multiple comparisons test. Significance levels: *$p < 0.05$, ns (not significant)

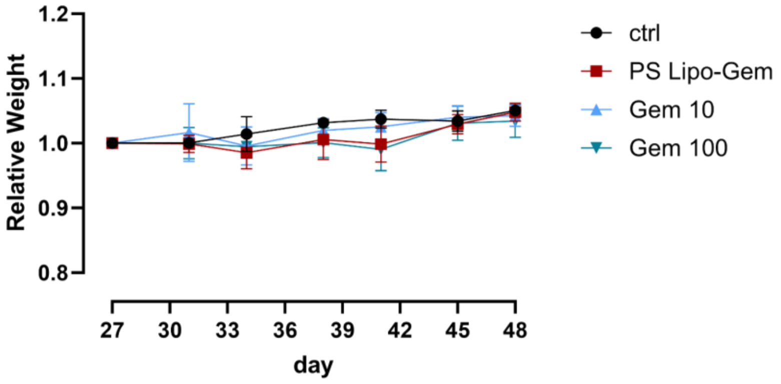


**Figure S8: Mice relative weight.** Throughout the experimental period, mouse body weights were systematically monitored. The weights were normalized to their initial value recorded on day 27, which marked the beginning of the injection phase. The treatments did not induce any discernible toxic effects, as the body weights of all mice were consistently increased. Data is presented as mean ± SD (4≤n≤6).

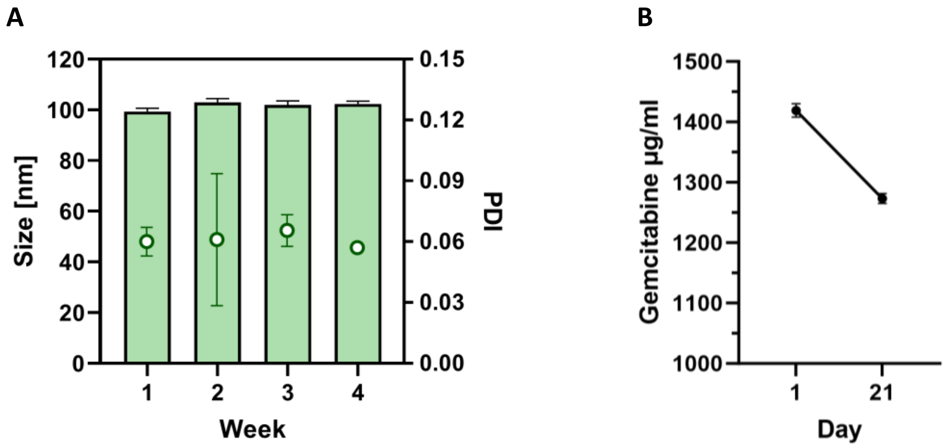


**Figure S9: The stability of injected PS liposomes. (A)** The size and PDI of gemcitabine-loaded PS liposomes were continuously monitored during the in-vivo experiment to ascertain their storage stability; **(B)** The gemcitabine encapsulation within PS liposomes was quantified on the initial day of encapsulation and one day after the final injection. Two biological repetitions performed in duplicates are represented as mean ± SD.

A

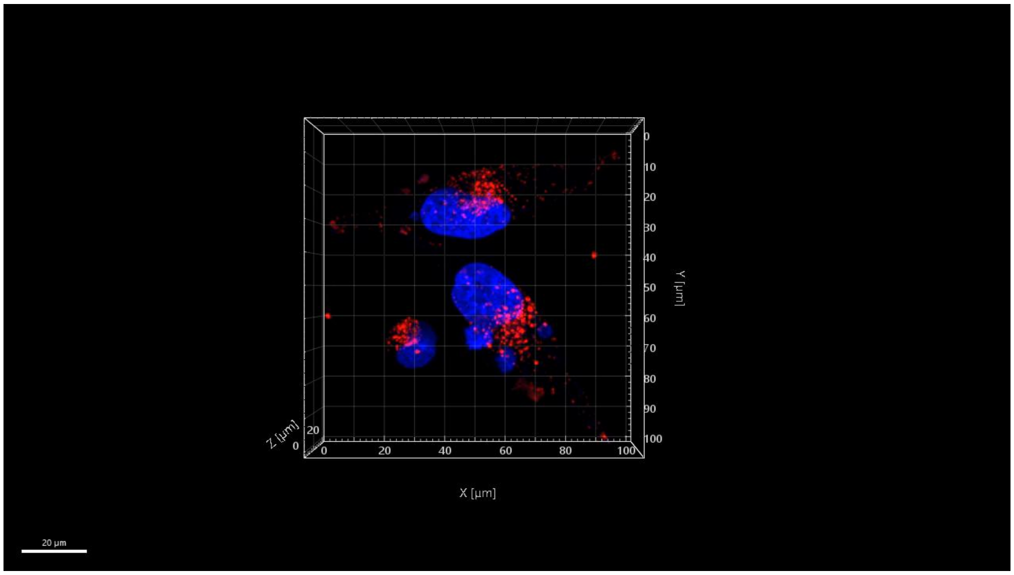


B

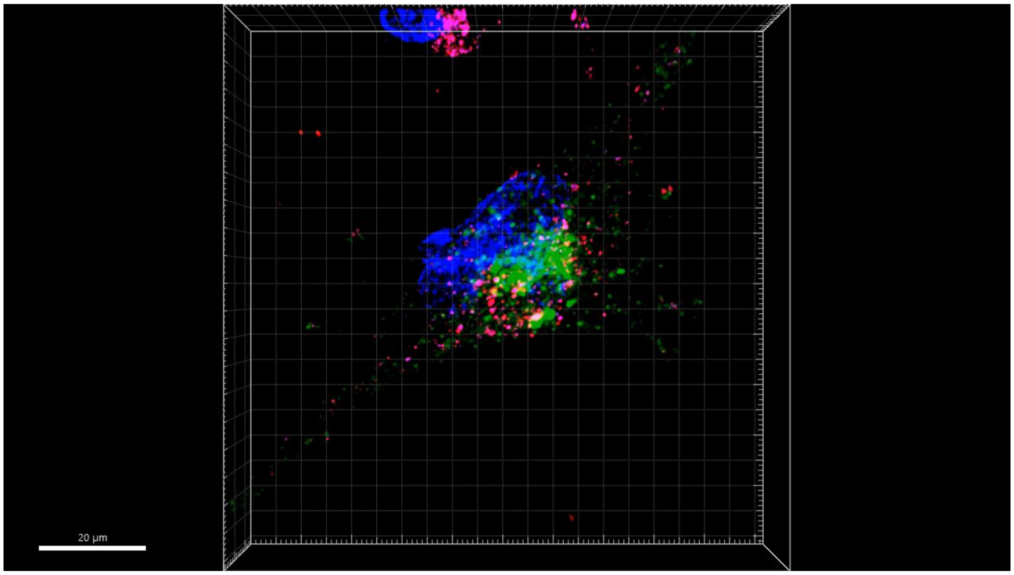


**Video S1: Nucleus and lysosome colocalization of PS liposomes.** Videos of 3D images of nucleus **(A)** and lysosome **(B)** colocalization with rhodamine-labelled PS liposome after 48 hours, produced by IMARIS; Scale bar 20 µm.
